# Earliest experience of a relatively rare sound but not a frequent sound causes long term changes in the adult auditory cortex

**DOI:** 10.1101/2019.12.24.887836

**Authors:** Muneshwar Mehra, Adarsh Mukesh, Sharba Bandyopadhyay

## Abstract

Sensory experience during a critical period alters sensory cortical responses and organization. We find that the earliest sound driven activity in the mouse auditory cortex (ACX) starts before ear-canal opening (ECO). Effects of auditory experience before ECO on ACX development are unknown. We find that mouse ACX subplate neurons (SPNs), crucial in thalamocortical maturation, respond to sounds before ECO showing oddball selectivity. Before ECO, SPNs are more selective to oddball sounds in auditory streams than thalamo-recipient layer 4 (L4) neurons and not after ECO. We hypothesize that SPNs’ oddball selectivity can direct development of L4 responses before ECO. Exposing mice before ECO with a rarely occurring tone in a stream of another tone occurring frequently leads to the strengthening of the adult cortical representation of the rare tone, but not that of the frequent tone. Results of control exposure experiments at multiple developmental windows and also with only a single tone corroborate the observations. A computational network model of known thalamic inputs to SPNs and L4 explains the observed developmental plasticity. Information-theoretic analysis with sparse coding assumptions also predicts the observations. Thus, salient low probability sounds in the earliest auditory environment cause long term changes in the ACX.

## Introduction

Sensory cortical circuitry development is activity-dependent and is based on the sensory environment to which an animal is exposed^1,2^. Even the earliest experiences, like in utero exposure to a mother’s voice, may lead to maternal speech preference in newborns^3^ and near term fetal responses to the mother’s spoken words^4^. It is well known that in the auditory system, altering the natural auditory environment during specific periods (critical period^5–7^) of development alters cortical circuitry and organization remarkably^8,9^. However, it is not known if the auditory system has the capability to adapt to environmental changes occurring before the known critical period. SPNs, the firstborn cortical neurons^10,11^, present in large numbers during and before the critical period^5,12^, play a crucial role in the development of functional cortical circuitry and organization^13–17^. Further, SPNs in the ACX are known to be the earliest to respond to sounds at least in ferrets^18^. However, the properties of early SPN sound responses, which are likely to play a crucial role in ACX development, are unknown, especially in the mice. SPNs receive inputs from the thalamic axons projecting to L4^10^ in the ACX^11,19^ and drive L4 activity over a time window of development^10,20^. Thus, it is important to know what kinds of natural stimuli specifically activate SPNs as their activity coincident with thalamic inputs can sculpt the earliest feedforward thalamocortical circuitry^13,14^ in the ACX. SP and L4 neurons play a crucial role in the maturation of thalamocortical circuitry, so it becomes important to know how the response properties of SPNs and L4 neurons change, starting from the onset of ACX sound driven activity.

Although ACX responses have been shown to exist in ferrets^18^ at ages equivalent to P6 (postnatal day 6) in mice^21^, no study reports the presence of sound driven activity in the ACX of mice at ages before ECO (P11/12). Low threshold hearing onset coincides with ECO^5,22^ in the mouse, with outer hair cell (OHC) based sensitivity allowing detection of near 0 dB SPL sounds. The developing auditory pathway is functional before ECO^23^ as known from ACX sound driven responses obtained by surgically opening the ear canal^24^ in mice. However, moderate-intensity sounds, for example, at conversational levels (~70 dB SPL) may drive spiking activity in the ACX before ECO without surgically opening the ear canal, through bone conduction^25^. The presence of responses before ECO suggests that activity-driven plasticity in the ACX starts earlier than previously thought. However, previous studies with a single tone exposure at ages before ECO, show no changes in tonotopy^5,7^ or ACX organization in the adult. Thus, even if responses in the ACX exist before ECO, it is important to understand the response properties in more naturalistic stimulus paradigms to test the possibility of activity driven ACX plasticity starting before ECO.

Thus, we first investigate the possibility of ACX responses of the L4 cortical plate and SPNs in mice with ear canals closed as would be required for the earliest natural sensory experience to drive cortical activity. With multiple techniques, we show that ACX sound-driven activity with ear canals closed start as early as postnatal day 7 (P7). We further show that, before ECO SPNs have unique response properties. SPNs, unlike cortical plate neurons, show selectivity to oddball sounds in a stream of repeating sounds before ECO. While after ECO, cortical plate L4 neurons show stronger deviant detection than remaining SPNs. The observed deviant detection shown by SPNs in the earliest phase can direct the thalamus to L4 synapse^13,14^. The above opens up the possibility that specific kinds of most initial sensory experience can modify cortical development. For example, exposure to music or the structured sound environment during the fetal period leads to long term plastic effects with enhanced responsiveness to the sounds experienced in the prenatal period^26^. We use a new auditory exposure paradigm with a low probability, salient/oddball tone (deviant, D) in a stream of a repeatedly occurring tone (standard, S). We show that with continuous exposure to the above S-D acoustic environment at ages before the established auditory critical period^5–7,27^ (before ECO), can remarkably alter the functional auditory cortical (ACX) responses, which persist into adulthood. The alterations are specific to the deviant or rare exposure tone (D). Performing control exposures with only a single tone during the same developmental window do not alter responses in the adult ACX. The observed plasticity is an outcome of the unique deviant detection properties we find of SPNs, before ECO, with strong responses to only the rare stimulus (D). With a computational network model derived from our experimental results and known thalamus to SP to L4 connectivity, we show how such plasticity occurs. Further, using mutual information maximization and sparse coding principles, we show that the outcome of early exposure with our new paradigm can be predicted theoretically. Hence our results revise the established timelines and concept of the auditory critical period and may generalize to other sensory systems.

## Results

### Mouse ACX and IC have auditory responses before ECO

We performed widefield fluorescence imaging of the ACX in Thy1-GCaMP6f (Jax Labs) at early ages (Fig. 1Ai) before (n=10; P7-P8=3, P09=4, P10=4) and after ECO (P13, Fig. 1Aii), to investigate the presence of sound driven Ca^2+^ responses in the ACX. After ECO, by P13, mice show strong Ca^2+^ responses and defined rostro-caudal gradient of high to low frequencies^28^ marking out the primary ACX (A1) and other auditory responsive regions (Fig. 1Aii). Since the auditory pathway is immature yet functional before ECO, we next investigate the presence of auditory responses to tones in the ACX of mice before ECO^5,22^ without surgically opening the ear canals. We found that sound-evoked Ca^2+^ responses are present in the mouse ACX at high (90 dB SPL) to moderate intensities (70 dB SPL, in awake state) at ages before ECO (Fig. 1Aiii-vi and Fig. 1B, Supplementary Video 1). With decreasing age, before ECO, progressively less percentage of areas of the cortical surface showed significant responses to multiple tone frequencies (Fig. 1Aiii-vi, Table S1), from 80% in P13 to 3% in P7. The percentage of frequencies of the tone stimuli tested, that showed significant responses in widefield imaging also progressively reduced (Table S1) from 71% in P13 to 20% in P7. Both the above reflect gradual maturation of the ACX starting from the onset of sound responsive activity at P7 to achieving tonotopy in A1 by P13. For further confirmation of sound-evoked responses in the auditory pathway before ECO widefield fluorescence imaging in the inferior colliculus (IC) of Thy1-GCaMP6f (Jax Labs) mice at P9-10 was performed. As in the ACX, we found significant sound-evoked responses in widefield fluorescence on the IC surface (Fig. 1C).

**Figure 1.**
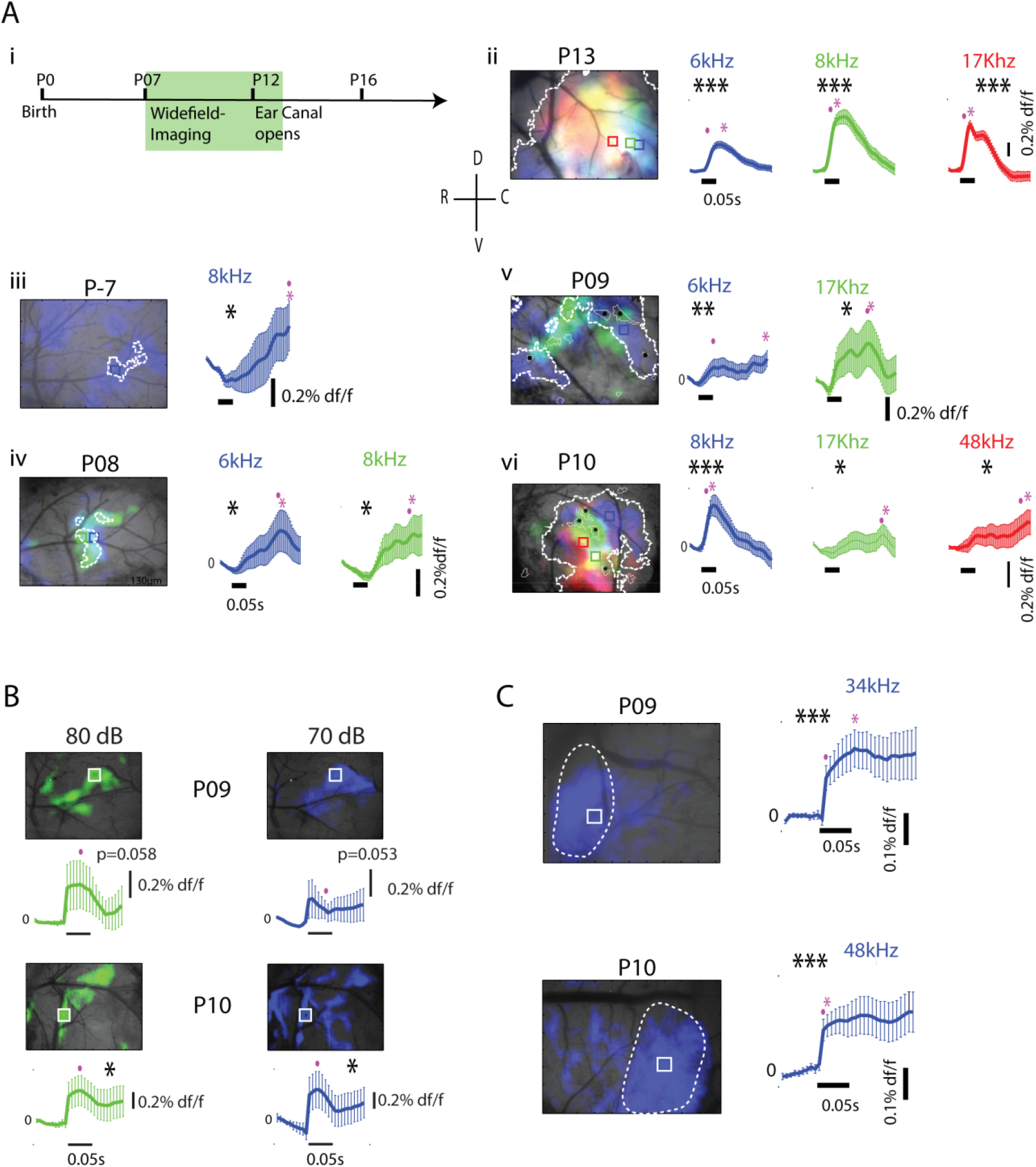
Widefield fluorescence imaging-based responses to tones in the ACX and IC of Gcamp6f mouse before ECO. A) i) Experimental timeline, ii) Widefield fluorescence imaging performed at P13 Gcamp6f mouse shows defined rostro-caudal gradient of high to low frequencies. Smoothed df/f traces with error bars representing SEMs of selected regions (square) responding to selected frequencies (color) are shown to the right of each image, iii-vi) Widefield fluorescence imaging-based significant response regions (white contours) in the ACX to 90 dB SPL tone in 4 different ages (P07-P10) before ECO. The black horizontal bar represents the stimulus duration. B) As in B, widefield fluorescence imaging in the ACX based significant response 80 and 70 dB SPL tone at P09 and P10 awake mouse. Respective df/f traces with SEMs of the selected region are shown at the bottom of each figure. C) As in A, widefield fluorescence responses before ECO in the IC at P9 and P10 to 90 dB SPL tone. The “.”(Purple) represents the first significant window, while the “*”(Purple) represents the most significant window. Sound-evoked Ca^2+^ responses are present in the mouse ACX at high (90 dB SPL) to moderate intensities (70 dB SPL) at ages before ECO (one-sided paired t-test, *p<0.05,**p<0.01 and ***p<0.001, Black).

### L2/3 and L4 ACX single neurons have responses before ECO

The presence of significant auditory responses in Ca^2+^ dependent fluorescence in widefield imaging (Fig. 1) could potentially indicate either the activity of input axons or spiking responses of, neurons in the ACX, or both. To rule out only input axon activity in the ACX widefield imaging, we performed 2-photon Ca^2+^ imaging of single neurons in L2/3 and L4 of the ACX of mice (14 mice) before ECO and after ECO (2 mice; P12-P13) (Fig. 2A) with high to moderate intensities of tones. Each 50 ms long tone stimulus was repeated 40 times and consisted of 3-5 frequencies from 6-48 kHz, ½ octave apart, presented at 60-90 dB SPL, with a 5-7s gap between repetitions. Single neurons in L2/3 in awake mice showed Ca^2+^ transients in response to tones at ages before ECO (P7-P9, Fig. 2Bi, Supplementary Video 2). The population mean df/f in response to different frequency tones (Fig. 2Bii) was investigated, considering all the neurons that responded with at least 25% reliability (fraction of repetitions showing a significant response, Methods). We found an increase in the mean population response strength before ECO (14 mice; P07-P08=3, P09=5, P10-P11=6, ANOVA with Tukey post hoc analysis, F(2,576) = 33.5, p <0.0001, Fig. 2Ci), and a decreasing trend in peak latency with age (Fig. 2Bii). The above indicated a gradual maturation of responses up to ECO (P12-13) in L2/3 ACX neurons. Along with an increase in tone response strength over age, response reliability increased significantly with age (Fig. 2Cii, 14 mice; P07-P08=3, P09=5, P10-P11=6, ANOVA with Tukey post hoc analysis, F(2,6953)=375.84, p<0.0001) before ECO, also showing gradual response development. The percentage of overall cases (imaged neuron and tone stimulus pairs) that showed significant responses also showed a gradual increase (Pearson correlation, r=0.76, n=6, p=0.078) with age from 6.84% at P7 to 12.5% at P12/13 (Table S2). All the above observations indicate the presence of sound driven Ca^2+^ activity before ECO in single neurons and that such activity varies with age in a manner expected from increasing maturity of the neural pathway.

**Figure 2.**
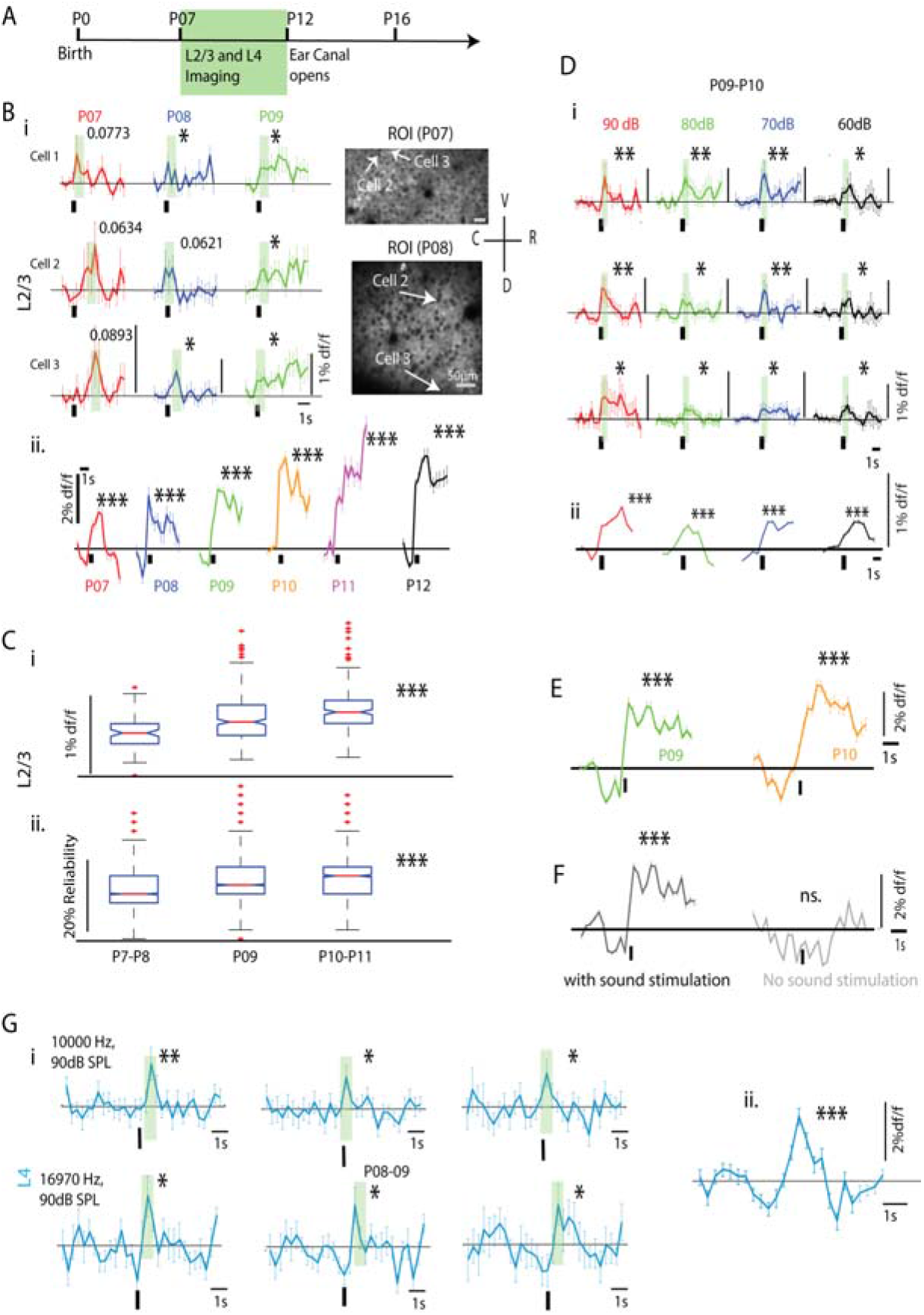
Sound evoked activity of ACX L2/3 and L4 neurons in the Gcamp6f mouse before ECO. A) Experimental timeline. B) i) Example traces of df/f of single L2/3 neurons (arrows in 2-P mean images of acquired frames with cells expressing GCamp6f at age P07 and P08, right) in response to 0 dB attenuation tones, respectively at three different ages (P07-P09). The black vertical line on the bottom of each trace represents stimulus onset. The corresponding green vertical shading bar in each example trace represents a significant moving window (3 frames) which shows the earliest significant p-value among all the frames (corresponding p-value shown above, one-sided paired t-test, *p<0.05). The error bars show the SEMs of 40 repetitions. B) ii) Population mean df/f of the age groups (color) of L2/3 neurons shown in Fig. Bi. C) i) Box Plot showing response peak strength of all L2/3 neurons imaged at different ages (14 mice; P07-P08=3, P09=5, P10-P11=6, ANOVA with Tukey post hoc analysis, corresponding significance-value shown to the right). C) ii) Box Plot showing response reliability of all L2/3 neurons imaged at different ages, as shown above (ANOVA with Tukey post hoc analysis, ***p<0.001, corresponding significance-value shown to the right). D) Similarly, as shown in B) i), are example traces of df/f of single L2/3 neurons of ACX in response to 90dB-60dB SPL tones at age group P9-P10 of an awake mouse, with corresponding significance values of the earliest significant window (3 frames) obtained, ii) The population mean df/f of L2/3 neurons across different sound intensity level (90dB-60dB SPL, 4 mice; 90 dB: 359, 80 dB: 348, 70 dB: 556, 60dB: 144 neurons). E) Extended 2-photon Ca^2+^ imaging data, showing population mean df/f (below) at two ages (P09, 2mice, 666 neurons and P10, 138 neurons) before ECO, indicating long (>2s) spontaneous window. F) Single neuron population mean df/f of the same age groups in response to sound (Left) and the same neurons without sound stimulation (Right), with 40 repetitions in each case (P09-P10, 3mice, 461 neurons). G) i) Three example traces of df/f of single L4 neurons (Cyan) in response to 0 dB attenuation tones at two different frequencies (Above row, 10 Khz and Below row, 17 kHz). The green patch represents the earliest significant moving window obtained with respective significance-values, ii) The population mean df/f of L4 neurons which showed significant responses (P9-10, 2 mice, 64 neurons). L2/3 and L4 ACX single neurons respond to sound before ECO (one-sided paired t-test, *p<0.05,**p<0.01 and ***p<0.001, Black asterisk).

The widefield Ca^2+^ responses at moderate intensities (Fig. 1B) were also ruled out to reflect only input axon activity as single neurons of L2/3 showed clear Ca^2+^ transients in response to tones down to 60 dB SPL (Fig. 2Di). The population mean df/f traces of all neurons (4 mice; 90 dB: 359, 80 dB: 348, 70 dB: 556, 60dB: 144 neurons) that responded significantly (with 25% reliability, Methods) to tones at the 4 intensities used, showed an increase in response strength with increasing intensity, except 80 dB SPL (Fig. 2Dii). The percentage of cases of single-neuron tone stimulus pairs that showed significant responses also showed a similar trend, increasing from 7% at 60 dB SPL to 40.5% at 90 dB SPL (Table S3).

The previous set of data were collected with usually a little over 1s of baseline preceding the stimulus. In order to rule out a contribution from any coincident spontaneous oscillating activity^29^ to the response, we obtained extended 2-photon imaging data before ECO (3 mice, P9 and P10, 6 ROIs) with >2s of baseline (Figs. 2E) and also with and without sound stimulation (Fig. 2F). The population mean df/f traces of all neurons in L2/3, showing significant activity above baseline in response to tones at P9 (666 neurons) and P10 (138 neurons) portray the same nature of responses as in Fig. 2Bii. The long duration mean baseline activity from 40 repetitions fluctuated around 0, as expected. We confirmed the presence of sound-evoked responses before ECO, considering any coincident ongoing spontaneous oscillations by comparing mean df/f traces of all neurons imaged in with and without sound cases (3 mice, 461 neurons, P9-P10, Fig. 2F). No significant activity was observed under no sound condition.

As in L2/3, putative L4 single neurons based on the depth of 2-photon imaging (300-350 microns from pia) also showed Ca^2+^ responses. Mean df/f based responses of 3 representative L4 neurons to 2 frequencies are shown in Fig. 2Gi. As in L2/3, the population mean of all significantly responding cases in L4 (2 mice, 64 neurons) showed responses to sound with >2s of baseline activity showing no spontaneous oscillatory activity (Fig. 2Gii). Thus based on 2-photon Ca^2+^ imaging of single neurons, we find both L2/3 and L4 to be tone sound responsive before ECO, showing responses as early as P7 and down to moderate intensities of 60 dB SPL. Together, the above data indicate that widefield fluorescence imaging-based sound responses in ACX are not due to only input axon related activity.

### SPNs in mice are driven by sounds before ECO

Two-photon Ca^2+^ imaging of single neurons, simultaneously from L4 and SPNs was beyond our scope^30^. Although SPNs are known to be the first to respond to sounds in ferrets, equivalent properties of SPNs in the mouse at the corresponding age of P6^18^ are unknown. Further, our Ca^2+^ responses do not show the nature of spiking activity in L4, thus requiring extracellular electrophysiology in mice before ECO. To reliably target SPNs with electrodes at different developmental ages, we first characterized the depth of SPNs based on SPN specific markers^31,32^ CTGF and Complexin 3 (Fig. 3B, green and red respectively) using IHC and based on cell morphology from Nissil stains^33^ (Fig. 3C). As the thickness of the cortex also varied with age, we quantified the SPN (Fig. S1E) and L4 (Fig. 3C, Fig. S1D) depth profile normalized to cortical thickness. The most consistent data was obtained with Nissil stains, and thus absolute depth as a function of age (P6 to P18) was used for electrode placements. Representative histology data are shown in Fig. S1A-E. The thickness of the SPN layer reduced with age as known^12,34^.

**Figure 3.**
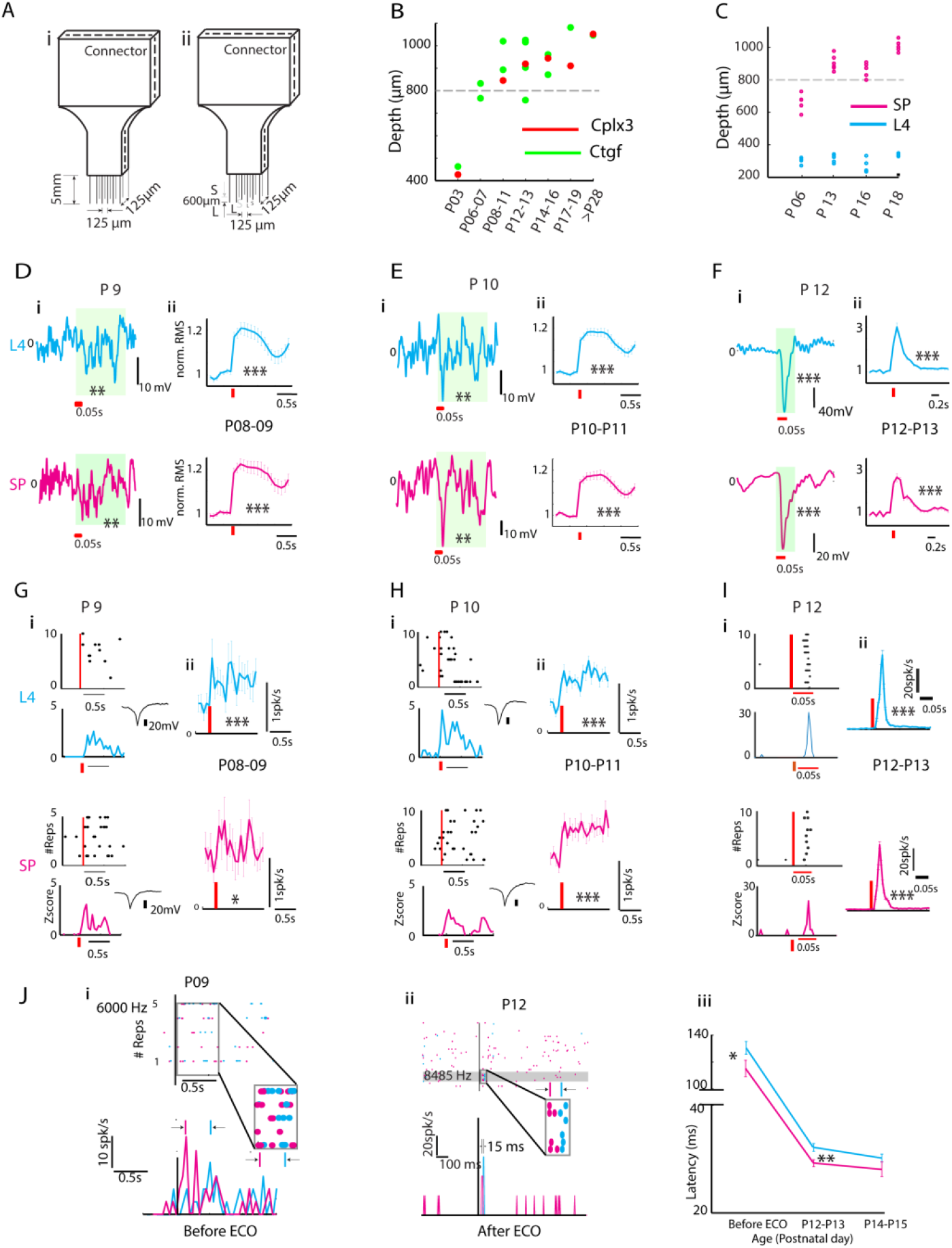
L4 and SP Neurons in the mouse ACX respond to tones before ECO. A) Schematic of types of 4×4 electrode arrays used – either all electrodes were of same lengths (i), or they had alternate short and long electrodes (ii) either 600 *μ*m apart or 400 *μ*m apart. B) SP depth determined with immunohistochemistry (IHC) by labelling SPNs with antibodies of SPN markers Complexin-3 (Red) and CTGF (Green). C) Depth of L4 (Cyan) and SP (Magenta) over age determined from cell morphology with Nissl stains. D-E) i) LFP in a single channel in L4 (Cyan, Above) and SP (Magenta, below), in response to tone at age, P09 and P10. The green patch represents the window across which the significance was calculated (L4 and SP ACX single neurons respond to sound before ECO (one-sided paired t-test, *p<0.05, **p<0.01 and ***p<0.001)., see methods). The horizontal red bar represents the stimulus onset. D-E) ii) Population average of all the significant recording sites and SEM of RMS values of LFP with corresponding p-values (one-sided paired t-test, ***p<0.001). The Red vertical line represents stimulus onset. F) i-ii) Similar Plots of single-site and mean population LFP activity at ages after ECO (P12-P13), as shown in D-E) i-ii), respectively. G-H) i) Dot raster and z-score of an L4 (Cyan, Above two Plots) and SP (Magenta, Bottom two Plots) neuron, in response to tone at age, P9 and P10. The Red vertical line represents stimulus onset. Inset to the right represents spike shape. G-H) ii) Population mean PSTHs of all the recorded SP (P8-9, 4 mice; P10-P11, 7mice) and L4 (P8-9, 4 mice; P10-P11, 9 mice) units across different age groups which had a z-score>1 in at least 1-time bin within 500ms after stimulus onset, with respective significance-values (one-sided paired t-test). The Red vertical line represents the stimulus onset. I) i-ii) Similar Plots of single-unit and mean population spike activity at ages after ECO (P12-P13) as shown in, G-H) i-ii), respectively. J) i) Latencies of L4 and SP neurons recorded simultaneously in P8-P11 (before ECO), ii) P12-13 and P14-15 age groups, iii) The population mean latencies revealed lowering of latencies with age and SPNs having lower latency than L4 neurons at ages before ECO and P12-P13 (one-sided unpaired t-test, *p<0.05 and **p<0.01).

We next performed extracellular recordings in the mouse before ECO (P7-P12) from the ACX SPNs and L4 simultaneously with custom made staggered electrodes arrays (Fig. 3A, 4*4 arrays x-y spacing 125 microns, 8 in SPN and 8 in L4, 400 or 600 microns apart in-depth, Microprobe, USA). Sound-evoked local field potentials (LFPs, Methods) simultaneously obtained from SP (>=800 microns; P8-P9, 3 mice, 72/536 and P10-P11, 4mice, 138/1248) and L4 (300-350 microns, P8-P9, 3 mice, 65/536 and P10-P11, 4 mice, 140/1248 locations) before ECO showed significant sound-driven activity (Figs. 3Di-ii, Figs. 3Ei-ii). Longer lasting oscillatory LFPs were observed at ages before ECO than after ECO (P12-P13, Figs. 3Fi-ii). The above results are in accordance with the pattern of LFP activity observed in ferrets at ages before ECO^18^. The population mean LFPs (Figs. 3D-Fii) were quantified with the root mean square (RMS) values in 50 ms bins normalized by the mean RMS values of the baseline (200 ms preceding the stimulus onset). Significant responses in both L4 (P8-P9; one-sided, paired t-test, t(64)=8.18, p<0.0001, P10-P11; t(139)=11.42 p<0.0001) and SP (P8-P9; t(71)=9.377, p<0.0001, P10-P11; t(137)=10.43 p<0.0001) were observed before ECO that last for > 1s (Figs. 3Di-ii and Figs. 3Ei-ii, Top and Bottom) unlike after ECO (Figs. 3Fi-ii, Top and Bottom, L4 and SP, respectively).

Single units in L4 (P8-P9, 4 mice; P10-P11, 9 mice, cyan) and SPN (P8-P9, 4 mice; P10-P11, 7mice, magenta) layer showed the presence of spiking responses to tones before ECO (Figs. 3Gi-ii and Figs. 3Hi-ii, Top and Bottom rows respectively) and after ECO (Figs. 3Ii-ii). Sample dot rasters (Top) and z-scores (Bottom) of spiking responses in 50ms time bins shown in Figs. 3Gi-ii and 3Hi-ii indicate the presence of significant sound-evoked spiking before ECO. Population PSTHs of all significantly responding units in SP (P8-P9; one-sided paired t-test, t(59)=1.733, p=0.043; P10-11, t(259)=10.07, p<0.0001) and L4 (P8-P9; SP, t(51)=5.738, p<0.0001; P10-P11, t(451)=12.678, p<0.0001) show significant sound driven activity before ECO, which get stronger with age (Figs. 3G-I ii). The response peak latencies (Methods) of L4 and SP single-units were obtained at ages before (P09 example trace, Fig. 3Ji) and after ECO (P12 example trace, Fig. 3Jii),. Before ECO and at P12-P13 the mean peak latencies of SPNs were found to be significantly lower than L4 neurons (one-sided t-test, before ECO, t(439)=−2.003, p=0.022; P12-P13, t(252)=−2.935, p=0.002, Fig. 3Jiii). The above results suggest that SPNs respond at shorter latency relative to L4 neurons before ECO, and the response properties mature with age. In subsets of experiments, at multiple ages, confirmation of electrode location in ACX was done with Dil crystal insertion in the recording site and observing labelling of MGBv (Fig. S1A). Also, in higher age groups, A1 recording was confirmed with the presence of general tonotopy (Fig. SIB) based on the spatial location of best frequencies (BFs) of units from multiple electrode penetrations. Electrode tracks were also routinely confirmed to be in the SPN layer (Fig. S1Di). In a subset of experiments in SCNN1-tdtomato mice (Jax 9613, P14, Fig. S1Dii) with staggered electrodes, confirmation of electrodes being simultaneously in L4 (L4 neurons expressing tdtomato) and SPN (from IHC, Fig. S1Dii) was done (3 mice, P12, P14 and P17). Thus based on the data from multiple techniques, we conclude that neurons in the different cortical layers and SP layer respond to sounds before ECO, without surgically opening the ear canal. Auditory responses are present as early as P7 in mice and respond down to 60 dB SPL tones.

### SPNs, unlike L4 neurons, show oddball selectivity before ECO

Having established the presence of auditory responses before ECO, we investigate the response properties of SPNs and L4 neurons in the context of stimulus-specific adaptation^35–37^ (SSA). We use oddball stimuli with a stream of standard tokens (S) with an embedded deviant token (D), to mimic the natural situation of a continuous auditory environment with important sounds occurring with low probability^35,37^ (Fig. 4A). The stimuli consisted of 50 ms long sound tokens presented at 3.3 or 4 Hz, of the pattern SSS…SDS…S, (D at 8^th^ or 7^th^ position of 15 or 10 total tokens) for the oddball response characterization. One of the S and D tokens was a tone *f*, range 6-34 kHz) and the other a broadband noise (*N*, bandwidth 6-48 kHz with equivalent sound level). Responses were collected in pairs with the S and D swapped to normalize for a neuron’s inherent selectivity for a particular S or D, (Fig. 4Ai). Each SD set of a pair was presented with 10-40 repetitions and 5-7s silence duration between the end of the previous and start of the next stream. A common selectivity index (CSI) was used to quantify a neuron’s deviant detection strength:

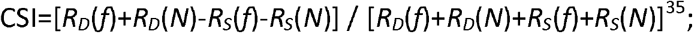

where *R_s_*(.) and *R_D_*(.) denote response to S and D, respectively. Two cases for *R_s_*(.) were considered: mean response to the first S or onset and mean response for all the S’s; the deviant detection strength values are referred to as CSI(F) and CSI respectively. To define oddball selectivity, if a neuron evoked higher responses to D compared to S in an oddball sequence with corresponding CSI value positive, then it was considered to exhibit oddball selectivity. We refer firing rate based oddball selectivity as deviant detection.

**Figure 4.**
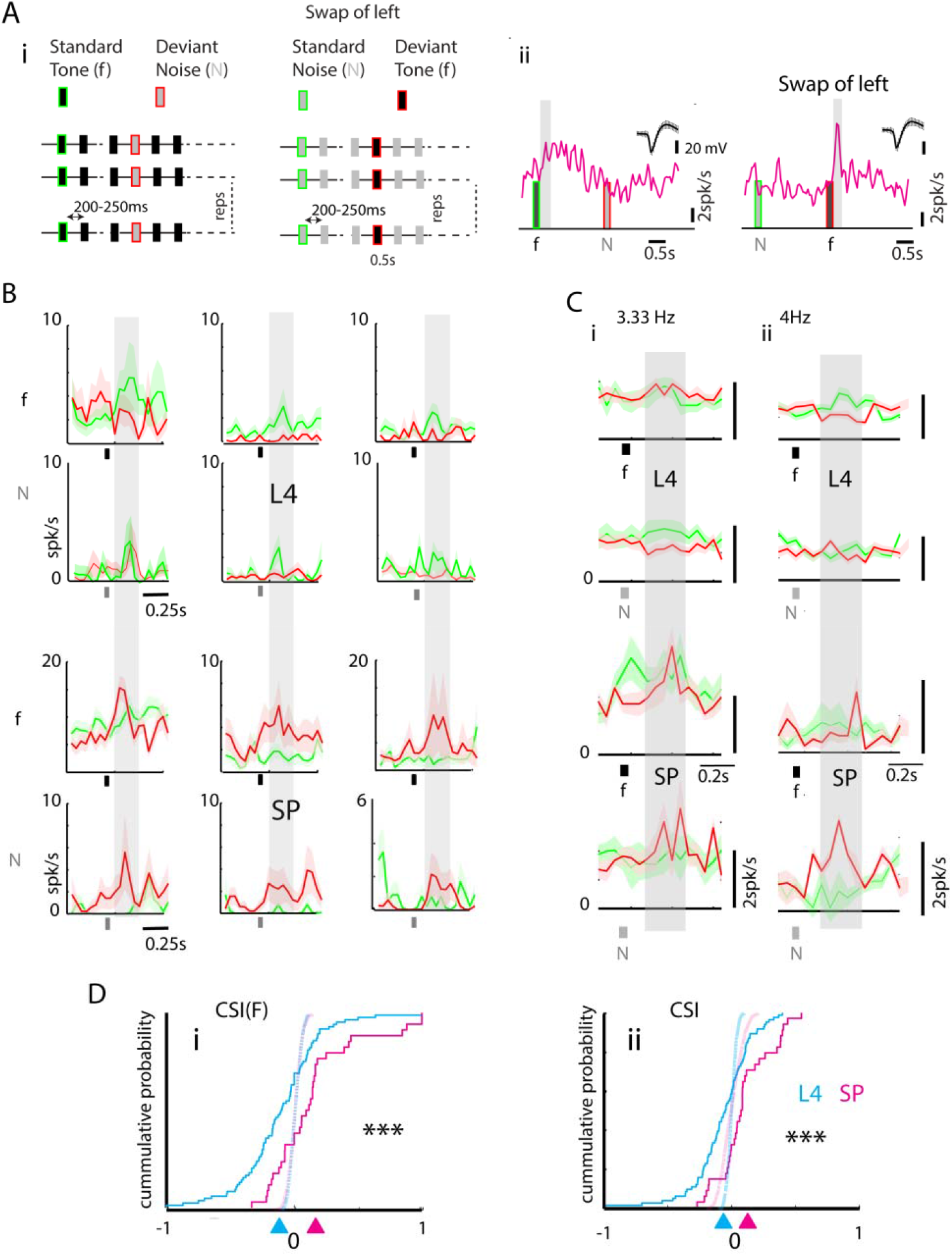
SPNs, unlike L4 neurons, show higher oddball selectivity before ECO. A) i) Schematic showing the Tone-Noise (f-N) oddball stimulation protocol with the swap on the right. A) ii) PSTHs of a single SP neuron, showing higher response at deviant tone (f) token (Swap, 250 ms response window gray patch, Right). B) Example traces of spike rate of 3 L4 neurons (upper 2 rows) and 3 SP neurons (lower 2 rows) in the first 3 columns, showing response to tone (f, Standard in Green and Deviant in Red, Above) and noise (N, Standard in Green and Deviant in Red, Below). A light gray patch is the mean rate window (250 ms) across which CSI(F) was computed. C) i) Mean population spike rate of all L4 and SP neurons (upper and lower two Plots, respectively) in response to tone (f, Standard in Green and Deviant in Red, Above) and noise (Standard in Green and Deviant in Red, below) presented at 3.33 Hz. C) ii) Same as C i), mean population spike rate of L4 and SP neurons in responses to tone and noise, delivered at 4Hz. D) i-ii) Population (3.3 and 4 Hz combined) Cumulative distribution of CSI(F) and CSI for SP and L4 (Magenta and Cyan trace, respectively) show significantly higher oddball selectivity in SP (one-sided unpaired t-test, ***p<0.001). Similar Cumulative distributions with mean CSI calculated for each of 1000 random pair of SD (Standard-Deviant) response windows selections for S response and D response either before deviant (Light dashed lines, Di) or after deviant (Light Dashed lines, Dii). No significant difference between L4 and SP in >90% of the random window sets.

ACX L4 (300-350 microns, 11 mice, 84 neurons) and SP (>800 microns, 11 mice, 34 neurons) single units responses were obtained before ECO in response to the presentation of the SD stimulus for *f-N* and its swap. Sample PSTHs of SPN before ECO shows the preference of the oddball or to the deviant tone (f) token (Fig. 4Aii). In general, response to the SD stimulus showed peaks in the response following the D-token, mainly in SPN and the first S-token or onset, primarily in L4 (Fig. 4B, L4 top row, SP bottom row). The earliest response peaks of L4 and SPN at these ages with the SD stimuli occur in a 250 ms window with a latency of 50-100 ms from the first token and in a similar window following the deviant (Fig. 4B). Comparing the mean response to the S-token and that of the D-token (Fig. 4B, S: green, D: red) shows the above. In the overall population mean response before ECO (P8-P12) for all L4 neurons, and all SPNs recorded indicate the presence of deviant detection in SP and not L4 (Fig. 4Ci-ii). Further, based on the spontaneous activity (100 ms preceding the stimulus), z-scores of the example neurons and population of neurons (Fig. S2) demonstrate a significant response to D and S and that in the population SP has significant activity at the deviant and not L4. To quantify the selectivity of the oddball, we obtaining spike rates in 250 ms windows because of the width of the peaks and calculated CSI(F) and CSI for all SPNs and L4 neurons. We observed higher deviant detection in SP than in L4 for both CSI (one-sided unpaired t-test, t(116)=3.666, p<0.001, Fig. 4Di) and CSI(F) (t(116)=3.954, p<0.001, Fig. 4Dii).

To rule out spurious deviant detection due to coincident spontaneous oscillating activity during the stimulus, we also considered CSIs expected by chance. 1000 random window pair sets each were considered in random-CSI calculations. Two non-overlapping windows 250 ms long were taken randomly in the standard region before the deviant or after the deviant not considering the deviant response. The mean CSIs were not significantly different between L4 and SP in >90% of the random window sets. More importantly, the distributions of mean random-CSIs for SPN and L4 showed that the means were not significantly different from 0 (dotted lines, Fig 4Di-ii).

Further to ensure that our results were not sensitive to the choice of latency and window size, we also varied the response window from 50 ms to 300 ms (in steps of 50 ms) and latency from 0 to 350 ms (in steps of 50 ms). We obtained the CSIs for all neurons, and we found for what combinations SPNs had significantly higher oddball selectivity than L4 neurons (Fig. S3). For a substantial range of values around the chosen latency and window size, the same results were obtained, indicating deviant detection in SPN and not in L4 before ECO.

### Unlike before ECO, L4 shows higher oddball selectivity than SP following ECO

Oddball selectivity in adults is a common phenomenon in all cortical layers of ACX^38^. Given the observed absence of oddball selectivity in populations of L4 neurons before ECO, we hypothesize that such selectivity in L4 starts after ECO. Thus, we next investigated how the oddball selectivity changes in L4 and SPNs after ECO. Single-unit responses to the same SD paradigm with *f*-*N* sound tokens (Fig. 4A) were collected simultaneously from L4 (P12-P13, 10 mice, 45 units, 57 cases; P14-P15, 8 mice, 66 units, 115 cases; P16-21, 4 mice, 20 units, 31 cases; P22-P28, n=6 mice, 38 units, 97 cases; >P28, 7 mice, 44 units, 63 cases) and SPNs after ECO from P12 to the juvenile/adult at >P28 (P12-P13, n=10 mice, 54 SPN units, 101 SPN cases; P14-P15, n=8 mice, 59 SPN units, 67 SPN cases). We again quantify the selectivity based on CSI(F) and CSI. Comparison of population PSTH segments in response to tone (Fig. 5Ai) or the noise (Fig. 5Aii) as the first S token (green), all S tokens (blue) and the deviant (red) show how selectivity to the oddball changes in L4 and SP over age after ECO starting from P12-13. Since direct comparisons are confounded by the inherent selectivity to the *f* or *N*, we compare CSIs over age (Fig. 5Aiii and iv) including the CSIs before ECO. L4 neurons start showing oddball selectivity immediately after ECO, which gradually declines to stable values with age till adulthood (CSI(F): ANOVA with Tukey post hoc analysis, F(4,358)=37.68, p<0.0001; CSI: F(4,358)=12.74, p<0.0001). On the other hand, SPNs with a significantly lower number at P12-P13 compared to peak at P8^34^, show significantly lower CSIs than L4 after ECO (CSI(F): one-sided unpaired t-test, P12-13_L4_ vs P12-13_SP_, t(156)=−1.944, p=0.027; P14-15_L4_ vs P14-15_SP_, t(180)=−1.10, p=0.13; CSI: P12-13_L4_ vs P12-13_SP_, t(156)=−1.621, p=0.050; P14-15_L4_ vs P14-15_SP_, t(180)=−1.41, p=0.07). We considered both CSI(F) and CSI since at early ages due to high adaptation (decrease in time constant with age: F_L4_(4,455)=11.92, p<0.0001) in neurons with robust responses at the first S token (responding tokens increases: F(4,455)=6.34, p<0.0001) and not any others (Fig. S4). The same trend was observed in both CSIs, showing that before ECO SPNs are stronger deviant detectors than L4 while after ECO the relative deviant detection strength switches.

**Figure 5.**
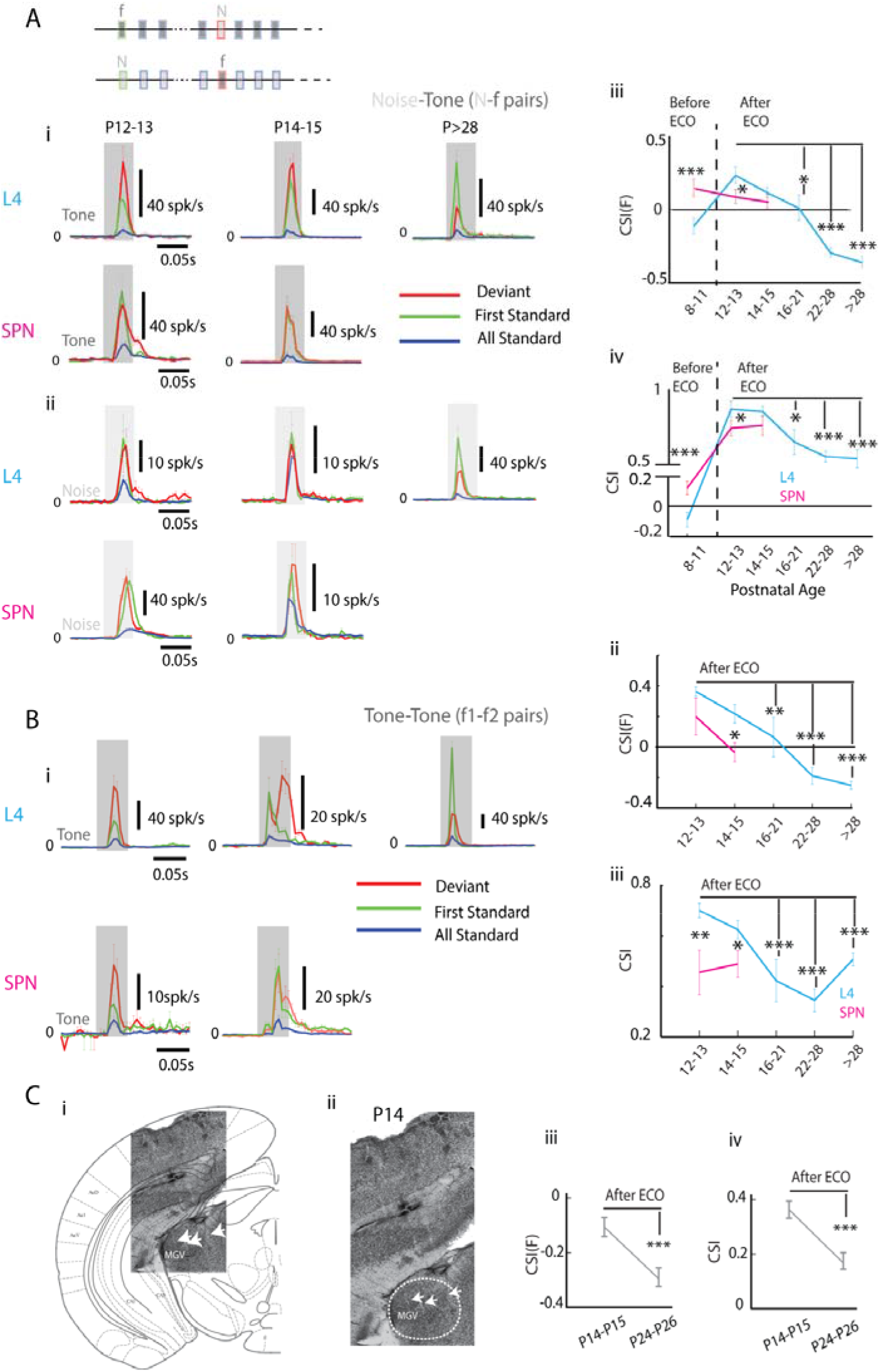
Sound evoked activity of ACX and MGB neurons after ECO. A) i) Schematic showing the Tone-Noise (f-N) oddball stimulation protocol with the swap, ii) Mean population PSTH of all significant responding neurons (L4 neurons above, SP neurons below), indicating the deviant response (Red), the first standard response (Green), and the mean of responses at all the standards (blue) to tone (Above 2 rows) and noise (Below, 2 rows) for different age groups after ECO. A) iii-iv) Overall mean oddball selectivity for noise and tone tokens measured by CSI(F) (above) and CSI (below), respectively as a function of age. B) i) Similar mean population PSTH for tone-tone (f1-f2) oddball stimulation protocol. B) ii-iii) As in A) iii-iv), Overall mean oddball selectivity for tone stimuli measured by CSI(F) and CSI respectively as a function of age. C) i) Example of electrode tracks (white arrowheads) shown in Nissl stains of brain slices showing recording locations in MGB of the thalamus, ii) Shows a magnified view of the relevant part of a coronal section with MGB, in which single-unit recordings from MGBv was performed (P14). iii) Mean and standard errors of CSI(F) (above) and CSI (below) for the thalamic neurons, by age groups, are shown, *p<0.05,**p<0.01 and ***p<0.001.

Above, we used *N-f* SD stimuli to mimic a repeating broadband auditory environment with the tone as a low probability stimulus (like a pup or adult mouse vocalizations^39^ in a normal auditory environment). Since deviant selectivity has been studied primarily with two tones^35,37^ (*f_1_* and *f_2_*), we also used an *f_1_-f_2_* SD stimulus set, to investigate if our results also hold in such a case. Further, using tones as both tokens allows more direct comparisons between responses. *f_1_* and *f_2_* were spaced either 1/4, 1/2 or 1 octave apart. Data were obtained from multiple L4 (P12-P13, 3 mice, 28 units, 47 cases; P14-P15, 3 mice, 41 units, 63 cases; P16-P21, 3 mice, 11 units, 15 cases; P22-P28, 4 mice, 24 units, 53 cases; >P28, 3 mice, 26 units, 74 cases) and SP (P12-P13, 3 mice, 9 SPN units, 14 SPN cases; P14-P15, 3 mice, 34 SPN units, 51 SPN cases) single units across different developmental windows. Data were collected from single units simultaneously, and hence *f_1_* and *f_2_* were at different distances from the BF of each of the units. One of the frequencies (*f_1_* or *f_2_*) was chosen to be at or near the BF of the majority of units. The responses to the SD paradigm were used to compare mean PSTH segments in response to the first S token, all other S tokens and the D token (Fig. 5B) as previously. A similar trend of decrease in CSIs in L4 over age was observed, as with *N-f* stimuli (CSI(F): ANOVA with Tukey post hoc analysis, F(4,247)=30.42, p<0.0001; CSI: F(4,247)=13.83, p<0.0001)(Fig. 5B). From CSI(F) and CSI, we found that L4 neurons are more selective (CSI(F): one-sided unpaired t-test, P12-13_L4_ vs P12-13_SP_, t(59)=1.8731, p=0.033; P14-15_L4_ vs P14-15_SP_, t(112)=2.866, p=0.002; CSI: P12-13_L4_ vs P12-13_SP_, t(59)=3.343, p<0.001; P14-15_L4_ vs P14-15_SP_, t(112)=2.163, p=0.016) to oddballs than SPNs after ECO.

The observed high oddball selectivity in SPN or L4 at different developmental ages could be a reflection of subcortical deviant selectivity^40,41^. The MGBv is the main thalamic nucleus from which A1, L4 and SPN get inputs^11,19^ (Fig. 5Ci-ii). Thus we quantified CSIs from responses of MGBv single units to the *N-f* SD stimuli (Fig. 5Ciii-iv). In both age groups considered (P14-15 and P24-26) we find that CSI(F) and CSI were lower than that of L4 (CSI(F): P14-15_L4_ vs P14-P15_MGBv_, t(180)=−4.373, p<0.0001; P22-28_L4_ vs P24-P26_MGBv_, t(163)=−0.031, p=0.488; CSI: P14-15_L4_ vs P14-P15_MGBv_, t(180)=−7.370, p<0.0001; P22-28_L4_ vs P24-P26_MGBv_, t(163)=−6.209, p<0.0001) and SPN (CSI(F): P14-15_SP_VS P14-P15_MGBv_: t(132)=−2.881, p=0.002; CSI: P14-15_SP_vs P14-P15_MGBv_: t(132)=−4.006, p<0.0001) in developing mice and also lower than L4 in the adult. Thus, the observed oddball selectivity even at the early ages is emergent in the cortex and expectedly a network phenomenon^42^.

### Exposure in a standard deviant acoustic environment before and after ECO show differential developmental plasticity for the standard and the deviant frequencies

Having found higher deviant detection in ACX SP than in L4 before ECO (P7-P11) and the opposite after ECO (P12-P15) during the known critical period, we next investigated the functional implications of our observations. Previous studies^5,7^ have shown that there is no change in the ACX organization in adult rodents following exposure to a continuous stream of a single frequency tone before ECO. Our results indicate that while L4 and SPNs are responsive before ECO, the above observation is expected given the strong SSA observed in SPNs before ECO (Fig. 4D), responding to the D stimulus and not the S. We hypothesize that because of higher deviant detection by SPNs before ECO (Fig. 4D), and because SPNs drive L4 neurons^19^ at early ages, exposure to an SD auditory environment before ECO could influence activity-driven plasticity of the ACX.

Adult ACX frequency representation of normally reared mice (EX0) was compared with mice exposed to an SD sound environment (S occurring with probability 0.9 and D with probability 0.1), at 4 different developmental stages between P6 and P21 (Fig. 6A, EX1-EX4). We exposed each of the groups to an f1-f2 combination of S=12 kHz and D=17 kHz. Additionally, to rule out frequency combination specific effects, for the most relevant age groups, namely P6-11 (EX2) and P11-16 (EX3), we also did additional experiments with another f1-f2 combination (P6-11: S=20 kHz and D=14 kHz, P11-16: S=14 kHz and D=10 kHz). In all the above cases stimuli at 90 dB SPL were used to ensure sound driven activity in ACX before ECO. Since we found that with ear closed, responses in the ACX existed for as low as 60 dB SPL, for our exposure study to be functionally relevant we did further experiments by exposing the P6-11 group at 70 dB SPL also (EX5). Finally, as stimulus control, we performed exposures of P6-11 (EX6) and P11-P16 (EX7) groups with an SS paradigm in which cases only one frequency (12 kHz) was used (EX6 and EX7 respectively).

**Figure 6.**
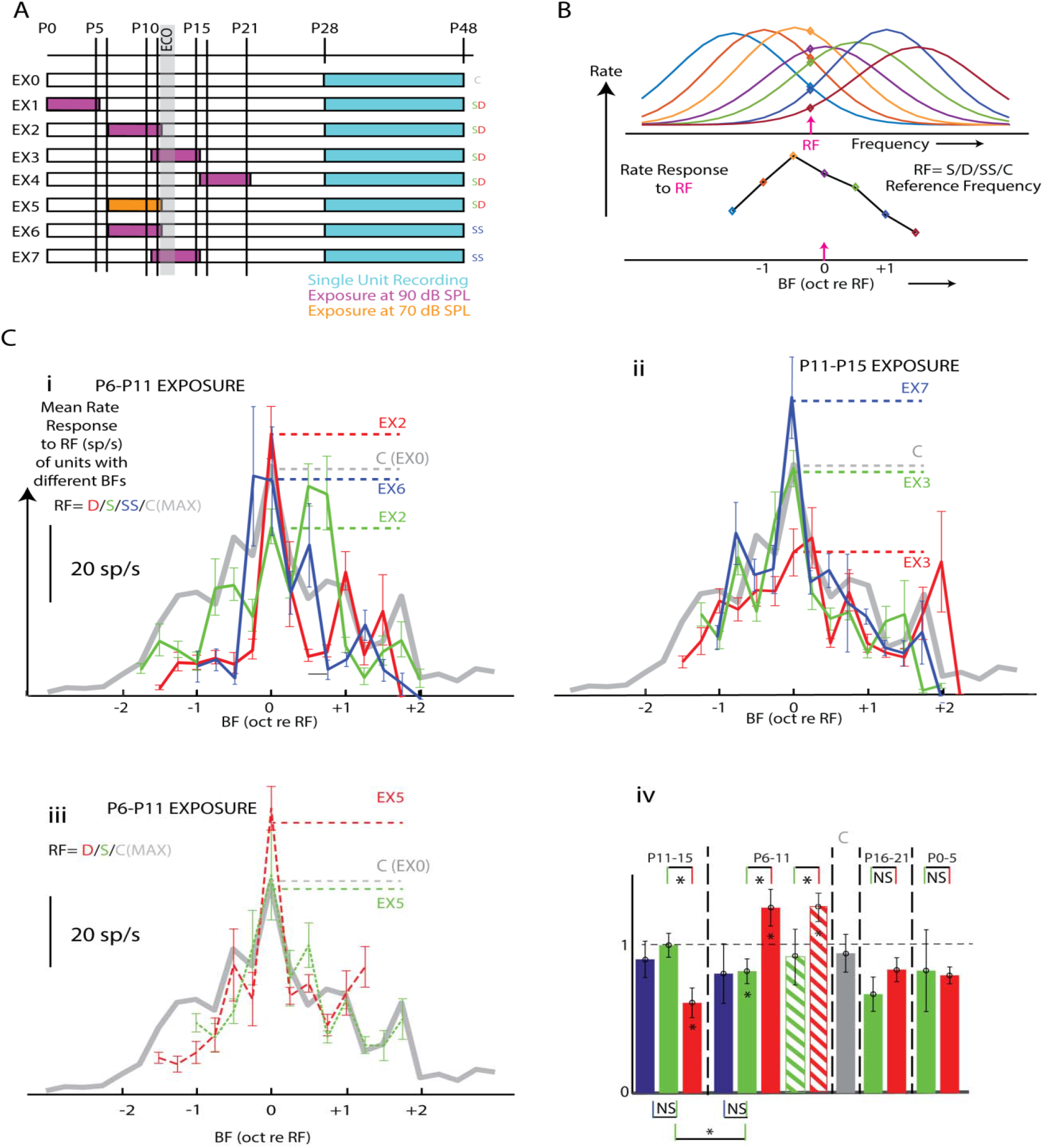
Exposure to an SD (Standard-Deviant) stimulus protocol, before ECO, induces long term plastic changes. A) Different exposure paradigms by age and type of exposure stimulus. B) Schematic explaining the construction of Reference frequency (RF) curves (see methods). C) i) RF curves of P6-P11 (before ECO) for SD and SS (only standard) exposure protocol at 90 dB SPL. ii) P11-P15 (after ECO), SD and SS protocol at 90 dB SPL, and iii) P6-P11 SD protocol at 70 dB SPL. The grey line represents the control group (EX0), showing the maximum boundary for all the 13 frequencies (see methods), iv) Bar Plots of all the exposure groups showing RF rates at BF normalized by the rates at all the other frequencies 0.5 octaves away on either side (see methods). The stripped bars in the P6-P11 group represent the SD exposure protocol at 70 dB SPL (EX5). The grey bar represents the control, *p<0.05,**p<0.01.

Single-unit recordings in L4 (depth 350-450 m) in response to pure tones (13 frequencies, 6-48 kHz, 1/8th octaves apart, 50-90 dB SPL, 10 dB steps) from A1 and AAF were performed in the exposed (EX1-EX7) and normally reared (EX0) mice at P28-42 (Fig. 6A). All significant response firing rates from the tuning curves (cartoon tuning curves, Fig. 6B) for each of EX0 to EX7 were obtained. EX1, EX4 and control (EX0) had the same median firing rate (rank-sum test, 12 sp/s for every case, p_EX1_=0.8475, p_EX4_,=0.1875). All other groups, EX2-3 and EX5-7 show, increased median firing rates (rank-sum test, 16-24 sp/s, p_EX2,3,5,6,7_ <0.0001 compared to EX0. Thus with auditory environment manipulations in the most relevant groups P6-P11 and P11-P16 (the known critical period), the overall median firing rates changed. With our limited resolution BF mapping, we did not find any apparent over-representation of any particular frequency in BF maps showing broad tonotopy (A1 and AAF) on the cortical surface (Fig. S5A) in any of the exposure cases, unlike other studies with the equivalent of the EX3 group (Fig. S5B). Other than the lower spatial resolution of single unit sampling, the above may also be due to parameters of exposure (sound frequency, duration and repetition rate). We considered only mice where recordings based on BF gradients were in A1/AAF.

Absence of over-representation of any particular frequency and increased firing rates in groups exposed during the known critical period and groups exposed during the period immediately preceding critical period, suggested plasticity in the strength of responses. Thus, we next investigated if the effect of exposure was on firing rate responses to tones and if changes were frequency specific. From the tuning curves, the population mean response rates to a reference frequency (RF) were Plotted as a function of the unit’s BF relative to the RF (Fig. 6B, Methods). The RF could be the frequency of S or D or that of SS. For S/D exposures (EX1-5) we obtained two such Plots, with S as RF and D as RF (green and red, respectively), for SS exposures (EX6-7), S was the RF (blue). For the control group C (gray) since there was no actual RF, each of the 13 frequencies presented was considered as RFs separately. Based on the 13 mean rate profiles as a function of BF re each of the 13 RFs, the maximum rate at each of the BFs (re RF) is considered as the upper boundary of observed rates at RF in control (EX0, Fig. 6C, gray), the lower bound being ‘0’.

The population mean rate as a function of BF relative to RF (re RF) for EX2&6, EX3&7 and EX5 (Fig. 6Ci, ii and iii respectively) showed differential effects on response strength at BF to the S and D for SD exposure before ECO and after ECO. Exposure after ECO in the known critical period of P11-15, the S frequency responses at S as BF are strengthened, in comparison to that of D (EX3; Fig. S6, one-sided unpaired t-test, t(107)=2.65, p=0.0045). However, before ECO, the opposite effect is observed (EX2; Fig. S6, one-sided unpaired t-test, t(120)=3.08, p=0.0013). No such changes are observed in exposure before P6 (EX1; Fig. S6, one-sided unpaired t-test, t(81)=0.15, p=0.55) and after P15(EX4; Fig.S6, one-sided unpaired t-test, t(58)=0.88, p=0.81). Similarly at 70 dB SPL exposure (EX5; Fig. S6, t(57)=1.507, p=0.06). Only in the case of SS for P11-15 exposure (EX7), there is an increase of S responses of neurons with S as BF beyond the boundary of control (gray dashed line, Fig. S6, one-sample t-test, t(17)=2.68, p=0.015). A higher response than the boundary of control for D was found before ECO SD exposure at both 90 (EX2, one-sample t-test, t(42)=2.80, p=0.007) and 70 dB SPL (EX5, one-sample t-test, t(38)=5.06, p<0.0001). Comparisons (significant or not) with a population of 13 control rate profiles are marked on respective bars (gray ‘*’ or NS). When considering each set of exposure frequencies, for the merged groups of SD exposure with different sets S and D frequencies, the response to D is greater than that of S before ECO (one-sided unpaired t-test, t(90)=3.07, p=0.0014, 20/14 kHz and t(28)=0.95, p=0.17, 12/17 kHz had comparatively less tuning curves, both 90 dB SPL and t(57)=1.5, p=0.06 12/17, 70 dB SPL). After ECO, the response for D was found to be less than that of S (t(51)=1.53 p=0.06 14/10 kHz and t(54)=2.11 p=0.019 12/17 kHz).

Absolute rates at BF for all frequencies could, in general, be higher after exposure. Although we did not get any over-representation, we did get a higher median rate for the relevant exposed groups. We investigated the effect of the exposure paradigm on the rates of all the neurons with respect to their BFs. To do that we made reference frequency Plots (Fig. 6B, see methods) and normalized the BF rates of the neurons with the BF as RF (center peak values of the Plots) by the mean BF rates of the neurons with BF >1/2 octave (two bins on either side) away from the exposed frequencies. A value more than 1 will signify a higher rate enhancement at the BF for the neurons with the BF as RF, while a lower value will indicate a higher rate enhancement at the BF for neurons with BF away from the RF. We found that in the SD exposure the contribution of the D frequency in the rate of neurons with D as BF is significantly higher before ECO (EX2) as compared to the contribution of S frequency in the rate of neurons with S as BF (Fig. 6Civ, bar Plots, one-sided unpaired t-test, t(120)=2.15, p=0.016 at 90 dB and t(57)=1.76, p=0.041 at 70 dB SPL) and the opposite is true after ECO (EX3, one-sided unpaired t-test, t(107)=2.71, p=0.004). The contribution of S in the rate of neurons with S as BF in the SD exposure before ECO (EX2) is lower than 1 (one-sample t-test, t(78)=−2.75, p=0.0078) but does not show any effect after ECO (EX3, one sample t-test t(75)=−0.83 p=0.39). Contribution of S at S as BF is not significantly different than 1 in the SS exposure both before (one-sample t-test t(6)=−0.37 p=0.72) and after ECO (t(17)=−1.23 p=0.23). As in the case of absolute rates, we looked into the frequency pairs individually and got similar results for before ECO (EX2, one-sided unpaired t-test, t(90)=2.16, p=0.016 20/14 kHz and t(28)=0.95, p=0.17 12/17 kHz had comparatively less tuning curves, both 90 dB SPL and t(57)=1.71, p=0.041 12/17, 70 dB SPL) and that of D is less than that of S after ECO (EX3, t(51)=1.61 p=0.056 14/10 kHz and t(54)=2.19 p=0.016 12/17 kHz). Thus there is a differential long term plasticity of responses to the low probability sound frequency depending on the period of development during which animals are exposed in the SD environment. It coincides with the differential oddball selectivity before and after ECO observed in the ACX L4 and SPNs involved early in development, in setting up the ACX circuitry.

### Model binary network with observed oddball selectivity of SPNs and L4 shows a stronger representation of deviant frequency before ECO and standard frequency after ECO

Having found long term plastic changes specific to the deviant in the ACX with exposure to an SD environment before ECO (P6-P11), we next explain our observations based on known circuitry of SPN, L4 and thalamic inputs^11,19^. SPNs receive stronger thalamic inputs than L4 early on and SPNs project to L4 neurons^19^ (Fig. 7A). In our simultaneous SP-L4 recordings with staggered electrodes before ECO, 42 pairs of neurons (between L4 and SP) were found to be connected (Fig. 7B-C and Methods) out of 136 simultaneously recorded L4-SP unit pairs. After ECO (P12-15), we found 176/244 L4-SP unit pairs to be connected. The direction of connections was obtained based on significant cross-correlations at positive or negative time lags and confirmed via granger-causality test. (SP->L4, magenta or L4->SP, cyan, corresponding examples in Fig. 7B and population data in Fig. 7C). Assuming columnar sizes of ~100μm, we grouped the pairs as columnar (≤ 125 μm) or not (>125 μm), based on the lateral distance between the electrodes from which connected pairs were obtained (Fig 7C). Observed directionality of the connected pairs before ECO suggests the existence of connections from SPN->L4 (12/16 pairs showed connections from SPN->L4 within a spatial distance of ≤ 125 μm, while 13/26 pairs were found to be connected at a distance of >125 μm, Fig 7C, first column, top). However, the L4->SP connection probability within ≤ 125 μm (4/16 connected pairs) was found to be significantly lower than SPN->L4 (one-tailed z test, z= −2.828, p=0.002, Fig 7C, first column, top). Moreover, L4->SP connections were found to be significantly higher than SP->L4 at ages after ECO (one-tailed z test, z=4.746, p<0.0001, Fig 7C, first column, bottom). The summary of connectivity between L4 neurons and SPNs (Fig. 7C) shows that SP->L4 connections are significantly reduced (one-tailed z-test, z= −2.600, p=0.004) from before ECO to after ECO. Granger causality (GC) analysis^43,44^ also revealed similar connectivity profiles between SP and L4 at ages before and after ECO, as observed using cross-correlation (Fig. 7C, second column, top & bottom). It is also evident from firing rates of L4 neurons that the thalamic drive increases after ECO as also observed in slice recordings^19^. We also found that recurrent connections, obtained with cross-correlation as well as GC, within SP (SP->SP) and within L4 (L4->L4) were differentially altered from before ECO to after ECO (Fig. 7D). Significantly more recurrent connections were present within SP than within L4 before ECO(z=5.437, p<0.0001, Fig. 7D, Above) and which switched after ECO (z=9.655, p<0.0001, Fig. 7D, Below). SP to SP recurrent connections decreased with ECO while that in L4 increased, very much like the observed CSI values in SPN and L4 before and after ECO, expected from the role played by recurrent connections in deviant detection^42^. Our observations on deviant detection strength measured by CSIs are thus further corroborated by the lateral connection changes observed in the two stages in the two layers.

**Figure 7.**
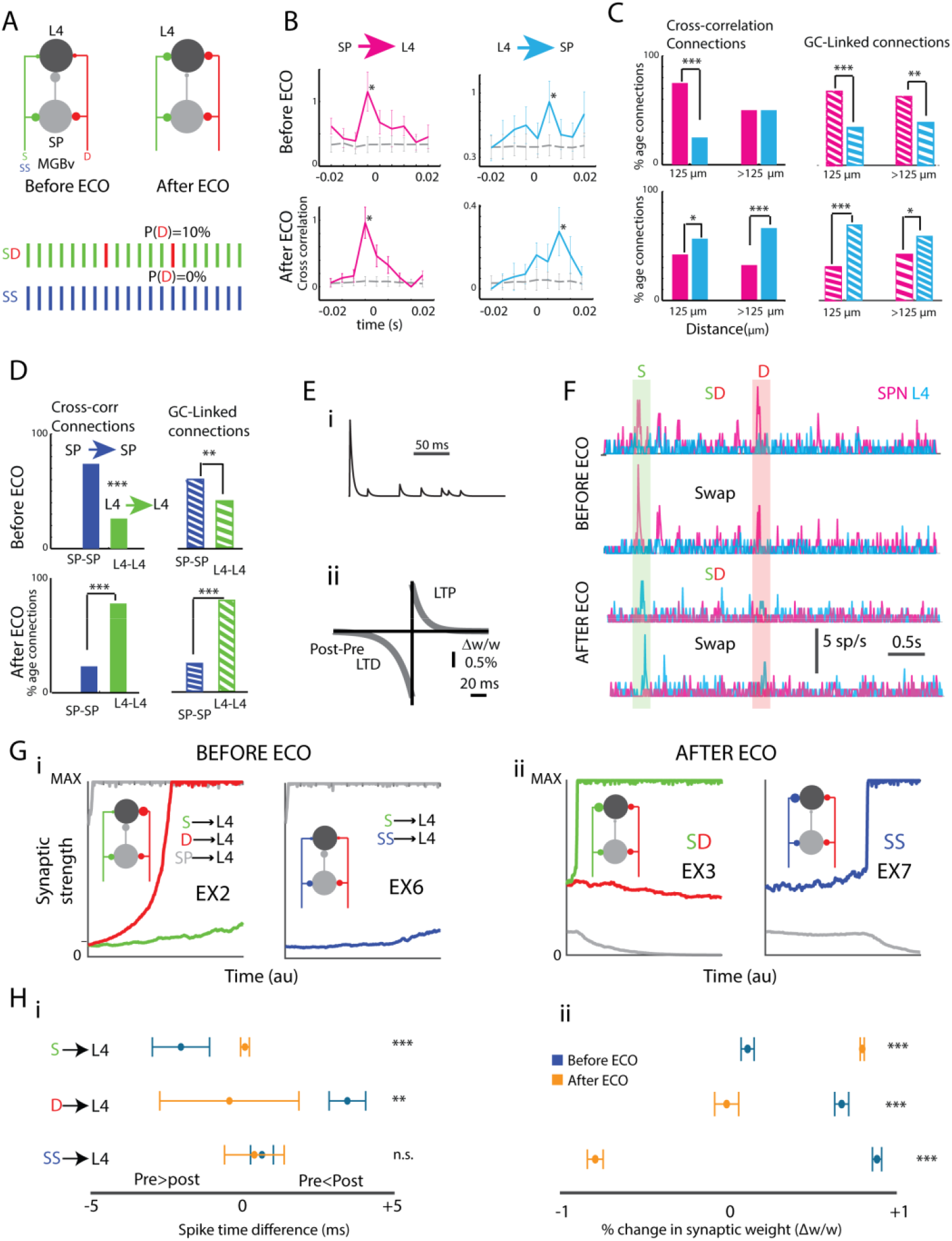
The binary network model of a developing ACX with subplate (SP) and thalamocortical (TC) inputs both before ECO, and after ECO, the size of the blobs represent the synaptic weight which changes across time. A) A network schematic showing standard (S, as Green in SD and as Blue in SS paradigm) and deviant (D, as Red in SD Paradigm) TC inputs to both subplate and layer IV and a subplate input to layer IV (top), Standard-Deviant (SD) and Only Standard (SS) stimulation protocol (bottom). B) Example pair of correlograms for SP to L4 (left) and L4 to SP (right) connections. The grey Plot represents the bootstrapped shuffled data (control). The “*” represents significant time windows (95% CI) for both before ECO and after ECO. An example EPSP profile, showing synaptic depression. C) Population bar Plots showing connection probability between SP and L4 for different distances according to cross-correlation(left) and Granger-causality (right). D) Connection probability for SP-SP and L4-L4 connections. E) i) An example EPSP profile, showing synaptic depression, ii) The asymmetric STDP learning followed by all the plastic synapses, showing a larger LTD window. The synaptic weight modifications occurred over −80 to +40ms of post-pre spike times. F) Example PSTHs showing higher oddball selectivity in subplate (Magenta) than layer IV (Cyan) before ECO and the opposite after ECO. G) Temporal evolution of the relevant synaptic strengths (S) in the two exposure periods (i: Before ECO, SD: EX2; SS: EX6 and ii: After ECO, SD: EX3; SS: EX7) for simulations of exposure with SD or SS protocol (the inset schematic shows the final point of the simulation with the blob sizes representing relative synaptic weights). H) i) Plots showing the mean spike time difference (post-pre) across time and ii) mean relative change in synaptic weight (Δw/w). *p<0.05,**p<0.01 and ***p<0.001.

Thus at the earliest stages stimuli through thalamic inputs that drive SPNs would drive the L4 neurons. Such activation of L4 neurons with coincident activity in the thalamus to L4 synapse would strengthen the synapse following Hebbian plasticity. Thus the observed deviant detection of the SPNs responding to D and not the S would strengthen the synapse on L4 conveying the D information. On the other hand, after ECO, the thalamus to L4 synapses are already stronger than before ECO, and the L4 neuron responds to both the S and D stimulus, preferring S more. The SP to L4 synapse is weaker than before ECO and is not as strong in evoking spikes in L4 by itself. Thus on exposure to SD after ECO, synapses conveying D information from the thalamus weaken, while that conveying S information strengthens.

We test the above explanation for the observed exposure dependent developmental plasticity based on L4 and SPN deviant detection through a binary network model (Supplementary Material, Fig. 7A) of early developing TC circuitry of the ACX^19^ similar to that considered in the visual cortex^14^. To have a minimal model that could potentially explain the results we did not consider lateral connections, although the presence of recurrence will, in principle, only improve the results. Thalamic inputs selective for either of two inputs, S or D (green and red, Fig. 7A) are considered to make synapses on an SPN, sending collateral inputs to L4 neuron^10^. The SP and L4 neurons are modelled as integrate and fire neurons with absolute and relative refractory^14^. The SPN neuron makes a synapse onto the L4 neuron as known^10^. All synapses were considered depressing in nature^42,45^ such that L4 and SPN responses show similar adaptation observed early in development (Fig. 7Ei and Fig. S4). Also, all synapses except the thalamic inputs to SPN are plastic following an asymmetric Hebbian spike-time-dependent plasticity (STDP^14^) rule (Fig. 7Eii). The abscissa in Fig. 7Eii represents the difference in pre-and post-synaptic spike times (*δτ*, post – pre), with positive *δτ* leading to LTP and negative *δτ* leading to LTD. The ordinate represents the percentage relative change in synaptic weight (Δw/w) for each pre and postsynaptic spike pair.

We consider the model with two different starting points, one representing the period before ECO (Fig. 7A, left) and another after ECO (Fig. 7A, right). In the earlier developmental period (before ECO), the SP CSI was 0.41, and that of L4 was −0.02, and in the later developmental period SP CSI was −0.17, and that of L4 was 0.4 (Fig. 7F), as in our observations (Figs. 5Aiii-iv). The differences in the initial point of the network (Supplementary Material) in the two cases (Fig. 7A left and right) are supported by other data^19^ and our observations of larger time constants later in development (Fig. S4), increased thalamic synaptic weights on L4 due to normal development, lower SPN to L4 connectivity (Fig. 7B-C) and reduced SPN rates through higher thresholds. We use the same exposure protocols SD and SS for the two periods to show the nature of how the thalamic inputs to L4 develop (Fig. 7G). Before ECO with higher oddball selectivity in SP than in L4, with an SD exposure, the D thalamic input to L4 strengthens faster than the S thalamic input (Fig. 7Gi). On the other hand, after ECO, with the oddball selectivity reversed in SP and L4, D to L4 synapse weakens and S to L4 strengthens, as L4 spikes more due to thalamic input spikes in S and not for that of D (Fig. 7Gii). The above is evident when considering the relative pre and postsynaptic spike times (δτ, post – pre) driving LTP or LTD of the S->L4 and D->L4 synapses in the two cases – before and after ECO (Fig. 7Hi, Supplementary Methods). The mean strength of weight change occurring in the synapses based on all Δw/w for each of the pre and postsynaptic spike pairs separated by δτ also show the same effect on synaptic strength change (Fig. 7Hii). Mean δτ(D->L4) is positive before ECO and negative after ECO with a significant difference. Thus the effect of SD exposure before ECO is overall long term potentiation of the D->L4 synapse while that after ECO the same synapse weakens as observed in the model results (Fig. 7G). The opposite is true for the S->L4 synapse. Before ECO with an SS exposure the S thalamic input does not show strengthening at a fast rate (as D, above) leading to the normal representation of the S input (Fig. 6Civ, bar Plot P6-11) equivalent to S in the SD case. After ECO, with the only S stimulus also the S inputs, expectedly strengthens leading to normal responses to S as observed (Fig. 6Civ, barPlot).

### Sparse coding and information maximization principles imply a strengthening of response to low probability stimuli

While a network model mechanistically addresses the observations, we theoretically address the observations of early exposure to a low probability stimulus leading to the strengthening of its responses and its weakening in later exposure case. Our observations and model results are contrary to the general ideas of long term plasticity - a repeated presentation of a stimulus in the critical period leads to the strengthening of the representation of that particular stimulus in the long term^1,5,7,9^. Auditory cortical spiking activity is known to be sparse and sparse coding principles theoretically explain several known receptive field types in the auditory pathway^46,47^. We assumed that a minimal activity-based coding in single neurons underlying sparse representation in populations and that such representation is one of the goals of the developing auditory system while maximizing information about the stimulus in the responses. With these principles, we use an objective function (Eqn. 1) as the desired optimization performed by activity-driven plasticity.

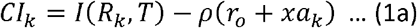

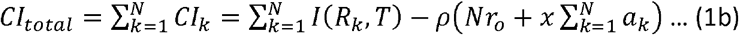

In Eq. 1, *I*(*R_k_,T*) is the mutual information (*MI*) between the stimulus (a random variable *T* which is either S or D, with probabilities (1-*x*) and *x*, where *x* is small and is considered as the probability of the deviant stimulus) and the response rate in the *k^th^* token is *R_k_, ρ* is a constant, *N* is the total number of stimulus tokens. *CI_k_* is considered as the constrained information function at the *k^th^* token. The mean ensemble response rates are assumed to be *r*_0_ and (*r*_0_ + *a_k_*) for S and D at the *k^th^* stimulus respectively, and assumed to have a Gaussian distribution both having variance *σ*^2^. The mathematical proof of the optimization is provided in the supplementary material. However, the basis is as follows. The mutual information would increase as the separation between the two rates (*a_k_*) increases. However, since the sparse activity requires minimal activity by single neurons to code the two stimuli, to minimize the summed activity, *a* must be positive and small. So the response to the deviant needs to be larger than that of the standard. However, the relative weightage of *MI* and the summed activity (*ρ*) can allow *a* to be increasing or decreasing and still cause *CI_k_* to be increasing.

We show that the condition for which *CI_total_* keeps increasing by considering Δ*CI* = *CI_n_* − *CI_m_* (*n>m*) with the assumption that the sequence {*a_k_*} is monotonic is

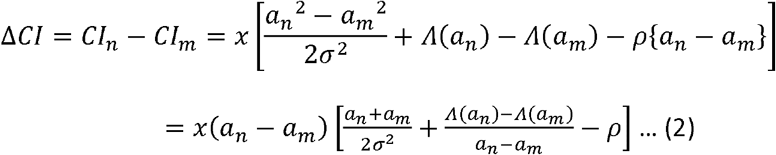

From the last expression in Eq. 2, under 2 different ranges of *ρ*, Δ*CI* is increasing with *an* increasing or *a* decreasing respectively. The theoretical analysis suggests that before and after ECO, the relative importance of coding or discrimination and minimal activity is changed. Based on the above, sparseness in later stages (after ECO), when spike rates are higher, needs to be weighted more than earlier stages (before ECO) to explain the differential effect observed before and after ECO in response to the low probability stimulus.

## Discussion

### Earliest sound driven activity in the mouse auditory pathway

We show that the mouse auditory pathway, ACX and IC, is driven by sounds with ear canals closed as early as P7 and down to intensities of 60 dB SPL. The above result is perhaps not surprising as auditory brainstem responses (ABR) in mice have been shown before ECO^23^ and that there are long term changes in prestin expression in mice when exposed to loud sounds at these early ages (P6-P11). Studies based on sound-evoked and spontaneous activity show that the central auditory pathway is functional before ECO. However, the peripheral auditory system may be immature at those ages. The external auditory meatus in mice reopens at P7, and the outer ear is fully mature by that time^49^. Further, lever arm lengths of middle ear bones are at 70% of mature values but have normal lever ratios by P7^50^. The oval window is also of normal size by P7^50^. Most importantly, mechano-transduction currents from outer hair cells (OHC) and inner HCs (IHC) become maximal in the cochlear base by P2 and at the apex by P5^51^’^52^, and thus IHCs are fully mature by P5^56^. However, OHCs are immature at P5 with nonlinear capacitance (NC) of OHCs starting at P7 and maturing fully at P18^53^. The development of OHC NC from P7 is correlated with the expression of prestin, the motor protein responsible for the OHC NC^53^ and hence hearing sensitivity. Thus, barring OHC NC, the key to low threshold (0 dB SPL) hearing and high auditory nerve fibre sensitivity, all evidence suggests that the peripheral auditory system is mature. Thus moderate to high-intensity sound-evoked activity in the auditory pathway is possible at ages before ECO in mice. However, no study to the best of our knowledge has shown auditory responses in mice before ECO. So far, all studies considering sound-evoked activity and early critical period in the auditory system have considered hearing onset to coincide with ECO. Thus our results open up an earlier time window of activity-driven plasticity to occur in the auditory pathway. Previous work on ferrets^18^ also shows such a possibility, where the authors show that SPNs are the first to respond to sounds at an equivalent developmental stage of ferrets as we find in mice.

### Auditory development and deviant detection

Post-ECO development of basic response properties like tuning, tuning width, thresholds to tonal sounds or broadband noise and other basic stimuli have been studied^5,54^. Our study not only shows that the starting point of auditory activity driven development is before ECO but also provides data on how deviant detection changes over development into adulthood. We show that the well-known deviant detection property of the auditory pathway^35,42^ is present in the early stages and is stronger in cortical neurons than the thalamus. We also show that deviant detection is strongest at early ages and reduces to the values observed in the adult. More importantly, we find that SPNs are deviant selective at the earliest stages of sound driven activity, before ECO, while cortical L4 neuron populations are not deviant selective at those ages. Since SSA underlying deviant selectivity is primarily a network phenomenon^42^, it may be surprising that SPNs are deviant selective at ages when thalamocortical and cortico-cortical connections are forming. However, it is well known that SPNs form synapses with each other^11,55,56^. Correlated with our observations of higher oddball selectivity in SPNs, SPNs are found to have more recurrent connections before ECO than after ECO and more than that of L4 neurons before ECO. After ECO, L4 neurons have more recurrent connections than before. The differential nature of recurrent connectivity in L4 and SPN before and after ECO also corroborates with our observations of differential oddball selectivity in the two populations, before and after ECO. Thus the nature of recurrent connectivity change we observe during early development supports the changes in the strength of deviant detection as measured by CSIs.

### Early auditory experience-based plasticity, SPNs and critical period

Broad tonotopic organization in the primary auditory cortex (A1)^57,58^ is known to be altered with over-representation of a particular frequency in the adult rats/mice which were exposed to that frequency tone during the critical period (P12 to P15, after ECO)^5,7^. In light of our present results with auditory responses present before the above period, we revisit exposure to a single frequency tone before the critical period. The previous studies^5,7^ did not find any evidence of adult cortical over-representation of the exposure frequency when exposing mice to that frequency before ECO. Thus, the window of P12-15 in mice and a similar window in rats was defined as the critical period^5,7^. Our data explain the lack of over-representation with exposure before ECO despite the presence of auditory responses during that period. The SPNs driving thalamocortical synapses to mature are found to have deviant detection at these early stages and thus SPNs, and consequently, L4 neurons do not respond to the repeated tone presentations. Therefore in our SS exposure results before ECO (P6-11, EX6) also, we do not find any plastic strengthening of responses to the S frequency as expected from above. However, the unique deviant detection properties of the SPNs before ECO, as hypothesized show strengthening of response to the deviant tone with an SD exposure before ECO. It is expected due to the lack of or weak responses to the repeating S frequency and responses to the D frequency. However, SD exposure following ECO shows the opposite since the SPNs are less deviant selective and thalamic inputs are now stronger, driving L4 responses directly. Similarly, the SS exposure after ECO also shows expected results. The mechanism underlying our observations is summarized in our model results. Of course, adaptation time constants play a role in the observed deviant detection and hence our exposure results.

Since SSA is primarily a network phenomenon with recurrent connections playing a key role, the observed changes in recurrent connections cannot be ruled out in understanding our results. In our current model, we did not consider the recurrent connections. Extension of our minimal binary network model to a recurrent network is expected to also produce the same results as the effects of deviant detection and SSA, underlying our observations would remain unchanged. Our model is only a minimal neural structure capturing the key mechanisms explaining the observations. Further, the model currently does not include inhibition, an element crucial in development, as synaptic plasticity rules of inputs/outputs of inhibitory neurons at such ages are not available. It likely plays a pivotal role as the maturation of inhibition is controlled by SPNs^14^. Also, the effect of the inhibitory switch, occurring before ECO, on the activity driven plasticity observed before ECO, needs to be investigated. The closure of critical period, around P16 also coincides with the observed near absence of SPNs in mice from P15. Thus mechanisms implicated in critical period closure like inhibition, adenosine, cholinergic inputs^6,59–61^, could also be tied to SPNs, since SPNs guide development of subcortical projections. Finally, exposures with selective inhibition of SPN activity during relevant periods of development are required to establish the SPN driven cortical development further.

We thus suggest that to understand experience-dependent developmental plasticity better, the period before ECO needs to be considered and possibly revise timelines of the critical period. The concept of the auditory critical period has been modified by showing that it can be stimulus-specific^62^. Similarly, we suggest the presence of a critical period that is deviant selective before ECO.

### Implications of the SPN deviant detection critical period

We show in mice that relatively low probability salient stimuli before ECO at ages equivalent to human gestational week 25^21^ cause long term plasticity. The above potentially explains observations in humans of maternal voice recognition in infants^4,63^. The results also suggest that prenatal sensory environment can influence cortical development and may also generalize to other sensory systems given the unique response properties of the SPNs. The generalization to other systems or other stimuli, in this case, is supported by our theoretical predictions. Although our results are derived based on tones as stimuli, the possibility of coding of auditory objects by the neurons in the ACX can also be addressed through similar experiments. For example, specific response selectivity to species-specific vocalizations in mice and other species could also be explained based on the low probability salient occurrence of such sounds during early development. Corresponding experiments for other sensory systems can lead to a better understanding of cortical development.

## Methods

All animal experiments were approved by the Institutional Animal Ethics Committee (IAEC) of Indian Institute of Technology Kharagpur. Detailed methods are provided in the Supplementary Materials. Thy1-Gcamp6f (Jax Labs) or C57BL6/J strain mice (NIN, Hyderabad, India or Jax Labs) across different developmental time windows were deeply anaesthetized for surgery (2-2.5% isoflurane), and prepared for in-vivo recordings after making a craniotomy (3-4 mm diameter, based on stereotaxic coordinates and landmarks). To investigate the presence of response before ECO, widefield and 2-Photon Ca^+2^ imaging was performed on Thy1-Gcamp6f expressing animal across different developmental window starting at ages before the ear canals were opened (P7 onwards). Few awake recordings were performed to probe responses at modest sound levels (70dB SPL). For in-vivo electrophysiological recordings from A1/AAF or MGB, 16 channel electrodes (Microprobes) were lowered in SP and L4 of ACX using a micromanipulator inside a sound chamber (IAC Acoustics). LFP and spiking responses to sounds (played through TDT free-field speakers ES1, attenuated by PA5 and generated with TDT RX6) were obtained with a unity gain headstage and preamps (Plexon Inc) and stored using National Instruments DAQ card. Sound generation, data acquisition and online/offline analysis were done with custom-written MATLAB routines. Anatomical labelling and confirmation of recordings from A1/AAF and from L4 or SP were done using multiple approaches (lipophilic tracer injections in recording site, Nissil sections and IHC). Exposure in specific auditory environments of mouse litters for the different time windows was done in another sound chamber (IAC Acoustics) maintaining an 8hr-16hr light-dark cycle. A computational model based on synaptic plasticity and fee-forward synaptic depression was added to demonstrate the high oddball selectivity before ECO. A mathematical commentary based on mutual information maximization was further done to provide theoretical support for our findings.

## Author contributions

MM performed all experiments, AM assisted. MM, AM and SB analyzed data. AM did modeling and theoretical derivation. SB supervised the work. SB, MM and AM wrote the manuscript.

## Acknowledgements

MM thanks CSIR for Fellowship, AM thanks MHRD for PMRF, SB thanks India Alliance, IIT Kharagpur, MHRD and SRIC Cell, IIT KGP for funding. This work was supported by the DBT/Wellcome Trust India Alliance Fellowship/Grant IA/I/11/2500270 awarded to SB.

## Supplementary Material

## Detailed Methods

### Subjects

All animal experiments were approved by the Institutional Animal Ethics Committee (IAEC) of Indian Institute of Technology Kharagpur.

### Acute craniotomy for wide-field imaging and 2-photon Ca^2+^ imaging

A standard surgical procedure was performed as mentioned above in in-vivo electrophysiology recordings. Animals were initially anaesthetized under isoflurane (induction at 5% isoflurane, followed by 2% isoflurane). After removing the skin and tissue above the skull, a metallic head plate was implanted above the estimated auditory cortical area. Craniotomy of around 3mm diameter was performed over the left auditory area. Smaller size craniotomy was preferred, as it gave us the advantage to minimize brain motion. A craniotomy was filled with 1.5% low melting-point Agarose. Immediately after that, a 5mm coverslip was imbedded over the craniotomy to dampen the pulsation. Animals were then transferred to the 2-photon imaging setup, and the level of anaesthesia was reduced down to 0.1-0.2% isoflurane during the recordings. For recording responses at lower sound levels, awake recordings were performed at age P9-P10. For awake recordings, Immediately after completing the surgery, pups were transferred to the sound chamber and allow them to recover from anaesthesia. This time window also allowed pups to habituate before the recordings were performed. Throughout the experiment, the animal temperature was maintained at 37°C by a heating pad.

### Wide-Field Imaging

For wide-field calcium imaging, imaging signals were acquired using a 14-bit CoolSNAP HQ2 CCD camera (Photometrics). Images of surface vasculature were acquired using blue LED illumination (470 nm), and wide-field GCaMP6f signals were recorded at ~30 Hz using blue illumination (470 nm)^1^. Fluorescence signals were acquired at 4x binning of 1040×1392 pixels, using a 4X objective (Olympus). Based on the auditory landmarks (Medial cerebral artery (MCA) and rhinal vein position) and the responses to different frequencies, the region of interest and the focal plane of the image (150-300 μm) were moved to desired coordinates for performing 2-photon imaging.

### Two-Photon Calcium Imaging

Two-photon calcium imaging was performed using a commercially available 2-photon microscopy system (Prairie View Technologies). The microscopy was controlled by Prairie View software (Prairie View Technologies). For imaging, the excitation beam of wavelength 860nm generated from Insight laser (Spectra-Physics, USA) was directed over the exposed brain volume. The laser was delivered through 20X/0.8 NA water immersion objective (2mm WD, Olympus). The laser power was adjusted from 50mW to 80mW depending upon the condition of the specimen. Frames in the region of interests (~150 μm x 200 μm with 1.16 μm pixel size) were imaged at ~4-6 Hz (160-250ms frame period, 1.2-4 μs dwell time) with the stimulus presented at usually the 6^th^ (to the 10^th^) frame in a sequence of 20-30 frames per stimulus.

### Auditory stimulation for imaging

The stimulus was presented through the electrostatic speaker (ES1) placed 2cm away from the right ear of the mouse pups Stimulus was generated through custom-written software in MATLAB (Mathworks), passed through TDT RZ6 multifunctional processor. Acoustic calibrations were performed using microphone 4939 (Brüel & Kjær, Denmark) and showed a typical flat (+/- 7 dB) calibration curve from 4-60 kHz. Auditory stimuli consisted of broadband noise (6-48 kHz, 40 repetitions), tones (40 repetitions of respective frequencies), and series of a sound token representing SD paradigm (30 repetitions). Only for one of the animals (P12, after ear canal opening), 5 repetitions of each pure frequency tone was delivered. Each of the stimuli was interleaved with a gap of 5s. The loudest intensities of sounds at 80-90 dB SPL were presented to ear, to allow mechanotransduction at younger ages (before ear canal opening). However, in awake recordings, 50-90 dB of sounds were presented at age P9-P10.

Single units spike times were obtained from the acquired spike channel data using threshold crossing and spike sorting with custom-written software in MATLAB.

### Anatomical labelling

Nissl staining was performed at different age groups to identify the SP layer and to guide in-vivo electrophysiology experiments. Mice of ages P6 (n=2 mice), P13 (n=2 mice), P16 (n=2 mice) and P18 (n=2 mice) were initially perfused, and the brains were harvested. Since ear canal opening is after P11 and as the cortical size stabilizes by P21, the above ages were chosen initially for the characterization. Nissl stains were performed after obtaining coronal sections of the auditory regions based on mouse atlas^2^. Immunohistochemistry was also performed on brain slices across different developmental time points (P06-P07, n=2 mice; P08-P11, n=3 mice, P12-13, n=5mice; P14-P16, n=2mice; P17-P19, n=2 mice; P20-28, N=3 mice; >P28, n=2 mice) with known markers of SP which are Complexin 3 (Cplx3); Synaptic Systems (122302) and Connective Tissue-Derived Growth Factor (CTGF); Santa Cruz (sc-14939). Paraformaldehyde (PFA) fixed brain tissue was embedded in a 3% agarose block. Brain embedded block was fixed on a vibratome (Leica) sectioning platform. 50 micron thin sections of auditory slices were cut, blocked and permeabilized in a buffer containing, 1% BSA, 0.3% Triton X and 10% Normal Donkey Serum (for CTGF) or 10% Normal Goat Serum (for Cplx3). Sections were incubated in primary antibodies (Cplx3: 1:500 and CTGF: 1:500) overnight and followed by washing with 1X PBS five times. After that, sections were incubated for two hours in Secondary antibodies (Cplx3: anti-rabbit 594, Invitrogen and CTGF: anti-goat 488, Invitrogen) diluted in the buffer, 1% BSA, 3% serum and 0.3% Triton X at room temperature. Sections were rinsed, mounted and imaged under epi-fluorescence microscopy. Depth of SP in different regions and at different developmental time scales was determined from these initial studies and was used as a guide for performing electrophysiology recordings. Further confirmation about recordings from input layer (Layer IV) was also obtained from SCNN1-td-tomato (cross of SCNN1-cre and floxed-td-tomato: JAX mice) mice, which have SCNN1 expressing neurons (specific to Layer IV) labelled, allowing confirmation of recordings in Layer IV. Sections were imaged using epi-fluorescence microscopy (Xcite illumination, Bruker), and were analyzed for cytoarchitecture by using custom-written codes in MATLAB.

### In-vivo extracellular single-unit recordings

Initially, animals were anaesthetized under deep anaesthesia (5% Isoflurane) inside the induction chamber. Initial Induction step helps in maintaining anaesthesia level in animals even after bringing down the anaesthesia levels (1.5-2%) during surgery. Core body temperature was continuously maintained at 37°C with a heating pad throughout the experiment. A central incision was made, and the left temporal portion of the skull was exposed. Exposed left temporalis was cleaned with the application of alcohol. After that, tissue clearance was done by rubbing the skull with 3% of Hydrogen peroxide. A 5mm diameter region was marked on the left hemisphere. Skull was firmly fixed after keeping the mark at the centre of a head plate. Craniotomy was performed over the estimated auditory cortical area. After completing a craniotomy, the experimental animal was shifted immediately to a soundproof chamber (IAC Acoustics). The depth of anaesthesia was continuously monitored throughout the experiment with paw pinches. During recording sessions, anaesthesia was brought down to a concentration of 0.5-1%, to get robust responses from primary auditory cortex.

Extracellular recordings were performed from the left auditory cortex using tungsten Micro Electrodes Array (MEA) of impedance 3-5 Mega ohms (MicroProbes, USA). A 4X4 custom-designed metal MEA with an inter-electrode spacing of 125 microns was lowered into the auditory cortex using a micromanipulator (MP-285, Sutter Instrument Company, Novato, CA). We also performed recording using arrays separated by 400 or 600 microns in length to record across laminae simultaneously (for example, Layer IV and SP, Fig. 3A). Signals were acquired after passing through a unity gain headstage (Plexon, HST16o25) and followed by PBX3 (Plexon) preamp with a gain of X1000. Wideband signal (used to extract LFP, 0.7 Hz to 6 kHz) and spike signals (150 Hz to 8 kHz) were obtained in parallel and acquired through National Instruments Data Acquisition Card (NI-PCI-6259) at 20 kHz sampling rate, controlled through custom-written MATLAB (Mathworks, Natick, USA) routines. Further, all online/offline analysis was performed using custom-written MATLAB routines. A laminar profile of responses was obtained from the primary auditory cortex (A1) by advancing the electrodes in depth at 100-150 micron steps.

Since the size and location of A1 over-development varies, it is essential to confirm the location of recording to be A1. For A1 recordings additional physiological information based on tuning – the existence of broad tonotopy, across electrodes in the recording array, latency of responses, selectivity to tone stimuli was used to confirm recordings in primary auditory areas (A1 and Anterior Auditory Field (AAF)). Based on the location of electrode tracks and comparison with mouse brain atlas^70^ and landmarks (location relative to the rhinal vein, size of the hippocampus in coronal slices) also provides additional confirmation of recording location. In many experiments after recording a Dil crystal^3^, a lipophilic tracer was inserted into the cortex at the recording location. After allowing passive travel of the tracer for 20-30 days at 35°C, brain slices were cut to look for labelled fibres and cell bodies in the ventral division of the MGB (lemniscal structure projecting to A1) allowing confirmation of recordings in A1 Fig. S1A). We further confirmed recordings from ACX by electrode location (using electrode tracks) in primary regions and also from broad tonotopy observed (Fig. S1B). The recording depths targeted for L4 and SPN were based on known depths of L4 and SPNs in ACX^34^; also, in a separate group of mice (P8-P11, n=3, P12-P13, n=7; P14-P16, n=5) depth of SPNs was determined to be above 800 μm based on two methods. First by cell morphology in Nissl stains of coronal slices (Fig. S1C) and second by immunostaining coronal slices containing A1 with antibodies of SPN markers^32^’^33^, *Complexin3* and *CTGF*, at different ages (Fig. S1E) and depth range of L4 from the expression of *SCNN1* (B6.Cg-Tg(Scnn1a-cre)3Aibs/J mouse, Jax Labs, 09613 crossed with CAG-tdTomato mouse Jax Labs, 07909), an L4 specific marker^34^. In a subset of mice, we confirmed electrode recording site to be in SP layer from electrode tracks along with immunostaining as above (Fig. S1E) and electrodes to be in L4 by performing recordings in the *SCNN1-cre-td-Tomato* (n=3, P12, P14 and P18) mice (Fig. S1D). Depths of other layers (L2/3 and L5/6) were determined based on the above results.

### Auditory stimulation for in-vivo electrophysiology

The stimulus was presented inside the soundproof chamber, 10 cm away from the right ear (contralateral) of the mouse, with TDT electrostatic speakers (ES1) driven by TDT drivers ED1 after attenuation by TDT attenuators (PA5) generated through TDT RX6 using custom made software written in MATLAB. The acoustic calibrations, performed with microphone 4939 (Brüel & Kjær, Denmark), of the ES1 speakers (TDT) in the sound chamber, showed a typical flat (+/- 7 dB) calibration curve from 4-60 kHz.

### Exposure protocol

For another set of experiment, animal litters were exposed to sound sequences inside a soundproof chamber with maintained 8-hr light and 16hr dark cycle, continuously for 5 days in 4 age groups (P0-P5/6, P6-P10/11, P11-P15/16 and P16-P21).

The litters were kept along with the breeders in a cage. Each exposure stimulus was synthesized online and presented through an electrostatic speaker, kept 5cm above the cage, using custom-written codes on MATLAB. All the age groups were presented with a sequence of stimuli having 50 ms sound tokens with a 250 ms inter-token gap at 70 and 90 dB SPL. The sequence (SD) had a token of two frequencies, a standard (S) having an arrival probability of 90% and a deviant (D) arriving with 10% probability. For only standard exposure (SS), there was no D frequency. After sound exposure, animals were returned immediately to home cages in the animal house and reared up in a normal environment until experiments were performed. During recording sessions, mapping of the primary auditory cortex was performed. Tone pips (50 ms, 5 ms up down ramps, 6-48 kHz, 0.25 octaves difference at 10-50 dB above threshold, 10 dB increments, 5 repetitions of each frequency-Intensity combination) was presented to the right ear. Extracellular responses were recorded from thalamo-recipient layer (Deep Layer III and layer IV) at depth 350-500 microns. Frequency responses area (FRA) was constructed of each unit for a minimum of three sound levels played. Single units from A1 having response peak latency within 9 ms to 31 ms were included in the analysis. This window was selected based on population mean and variance of response peak latencies obtained across all sound levels.

### Data Analysis

#### Wide Field imaging analysis (Fig. 1)

Wide Field imaging analysis was performed using custom codes written in MATLAB (Mathworks). For analysis of the wide-field data, raw images collected were binned at 4x and then smoothed with a moving window of 4×4. Further analysis was done on the smoothed images. To find the significant pixels, a one-sided t-test across all the iterations was performed between the 5 frames immediately preceding the stimulus onset and a moving window of 5 frames after the stimulus onset in all 130 μm by 130 μm square areas. The pixels which gave a p-value of less than 0.05 on at least one window after the stimulus onset, were considered for further analysis. The mean change in df/f with time traces was constructed for pixels showing significant positive responses. These traces were further smoothed with a moving average window of 5 frames, and a one-sided t-test was performed again to find the first significant post-stimulus frame and the frame with the most significant response.

**Table S1.**
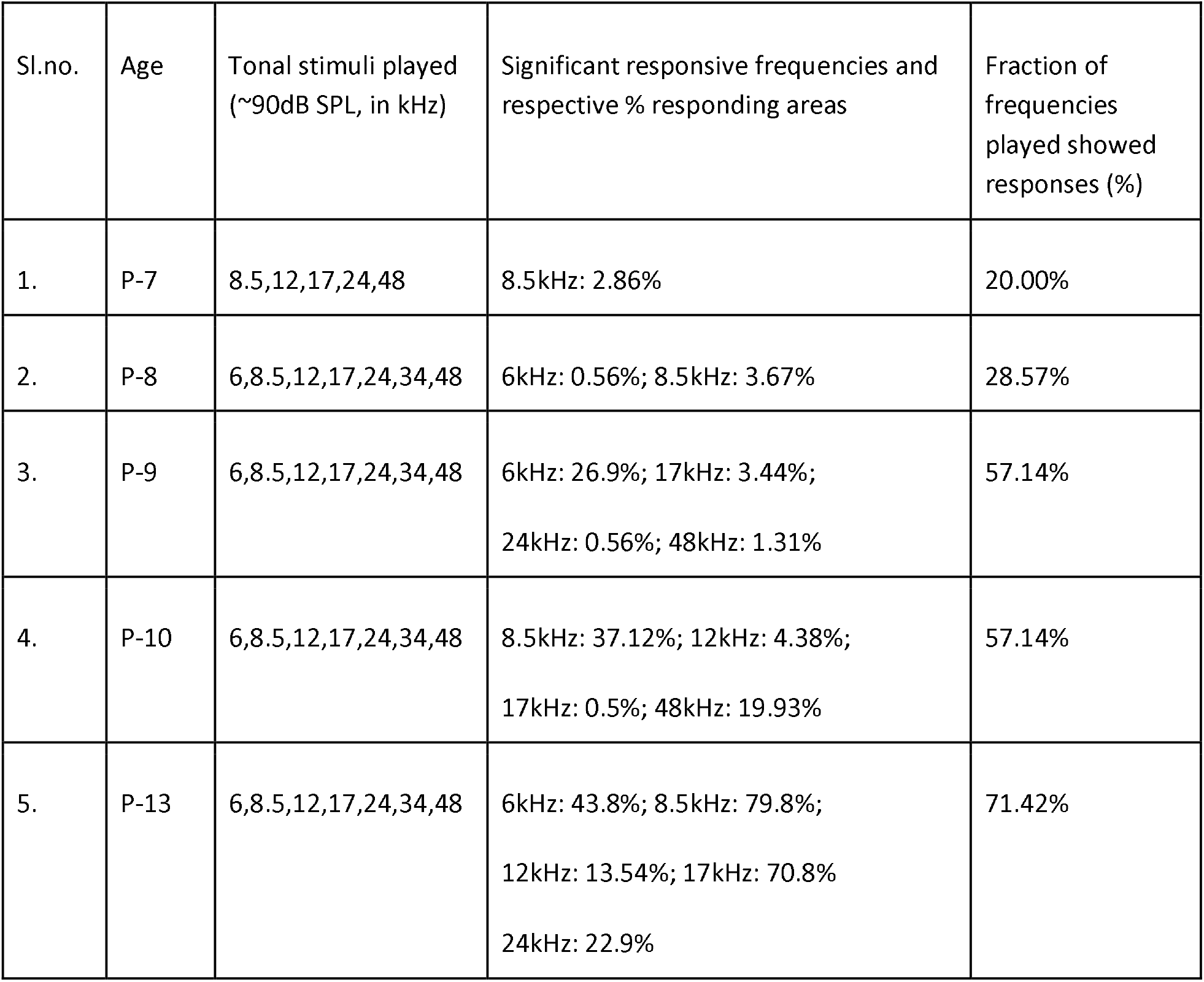

#### 2-photon imaging analysis (Fig. 2)

2-photon imaging analysis was performed using custom codes written in MATLAB (Mathworks). Imaging sequences were aligned by performing X-Y drift correction. Cells were selected manually by selecting the centre point of the cell on the motion-corrected mean image. ROI (5 μm radius) were drawn based on the cell centres. Raw fluorescence signal over time (F) of the selected ROIs across all frames were extracted. For each trial, relative fluorescence was computed by using ΔF/F_0_ = (F − F_0_)/ F_0_), where F_0_ corresponds to baseline fluorescence. Baseline fluorescence amplitude was estimated by calculating the mean of fluorescence values over all the frames preceding stimulus, except the first 2 frames. Neurons on stimulation to tones and noise were considered to be significantly responsive as follows. One-sided paired t-tests were performed between mean df/f over 3 of the successive frames before the stimulus and each one of 6 windows, of 3 consecutive frames after stimulus onset (up to the 8^th^ frame, ~1.5-1.8s). If any of the 6 moving windows showed a significant increase in mean df/f (40 repetitions) over baseline, the neuron was considered responsive.

**Table S2.**

| Sl.no. | Age | Fraction of cases (all neurons in response to different frequency tone at 0 dB attenuation) that showed significant responses. |
| --- | --- | --- |
| 1. | P-7 | 6.84% |
| 2. | P-8 | 5.6% |
| 3. | P-9 | 10.6% |
| 4. | P-10 | 15.4% |
| 5. | P-11 | 12.3% |
| 6. | P12-P13 | 12.4% |

**Table S3.**

| Sl.no. | Tone Intensity | Fraction of cases (all neurons in response to different intensities of tones) that showed significant responses. |
| --- | --- | --- |
| 1. | 90 dB | 40.50% |
| 2. | 80 dB | 12.77% |
| 3. | 70 dB | 33.04% |
| 4 | 60 dB | 7.02% |

For significant responses in single trials to compute reliability, a trial was deemed responsive if the mean df/f of 4 frames (~1s) after the stimulus after stimulus onset was higher than 1*STD (STD obtained from 40 repetitions) of the mean of the 4 baseline frames preceding the stimulus. Population analysis was performed on only those neurons which evoked significant responses to >25% of the trials (>25% Reliability). Reliability was computed as the proportion of trials that showed significant responses out of 40 trials.

#### Identification of Subplate Layer (Fig. 3, Fig. S1)

The identification of the SP Layer was based on separation drawn by sparse cell zone above, and the horizontally oriented cytoarchitecture of the SP neurons. Based on the above features, manual points (perpendicular to lamina) using MATLAB on slice images were selected to define the depth and width of the SP layer. A similar analysis was performed too on the immunolabelled brain slice images.

#### LFP analysis (Fig. 3D-F)

For LFP analysis, baseline shifting was performed to have a mean of baseline 200 ms preceding the stimulus at 0. To obtain LFP, acquired wideband signals were notch filtered (50Hz, Butterworth 8th order, to remove AC supply line noise) and then band-pass filtered (between 5 Hz and 300 Hz, Butterworth 2nd order). LFPs were smoothed further using a Gaussian window, SD 150 ms. LFP responses crossing the threshold of +/- 200 mV throughout the recorded time window were excluded from population analysis. A spontaneous window of 400ms before the onset and a response window of 500ms after the onset was considered for testing the significance of LFP response. The entire stretch of 900ms (spontaneous + response) was divided into 9 bins of 100ms each, making 4 bins in the spontaneous window and 5 bins in the response window. One-sided paired t-test was performed between the mean of the four spontaneous bins and each of the five response bins and only those traces with a p-value less than 0.05 in at least one of the five response bins were taken for further analysis. The normalized population RMS Plots were Plotted for these traces, where each of the post-onset RMS values was normalized with the mean of the 4 spontaneous RMS values.

#### Rate Calculation

Neuronal firing rate was computed by calculating the mean response within stimulus duration or token. Only single units that responded (compared to spontaneous activity 400ms preceding stimulus, 2-tailed t-test, p<0.05) to at least one of the stimuli (sound token (*f* or *N*) as standard or deviant) used were considered for analysis.

#### Latency Analysis (Fig. 3J)

For each neuron, tuning parameters (Best Frequency (BF) and Latency) were defined using custom-written codes in MATLAB. The best frequency for a neuron after ECO is the frequency which elicits a maximum response within stimulus duration (50 ms) to a given frequency at a given sound level. At ages after ECO, Latency to stimulus onset was defined as the time bin (5 ms resolution) at which peak response was observed for a BF. For calculating latency at ages before ECO, we first considered all Psths of neurons binned at 5ms with z score >1 within 500ms after stimulus onset. After that, a peak response within 300ms after stimulus onset was selected as the latency to the presented stimulus.

#### Adaptation Time Constants (Fig. S4)

A time-course analysis was performed to fit the response rates for SSS…..SDS….SS stimuli. Across all age groups and cortical layers, we emP10yed exponential fitting to only those responsive units which have significant responses for tone as standards. To compute exponential decay time constants, the average spike rate in each token before the deviant was fitted with a first-degree exponential curve, based on least square error method. Fitting was performed using the ‘fit’ function in MATLAB.

#### Construction of Reference frequency Plots (Fig. 6)

To investigate if the effect of exposure on firing rate were frequency-specific, the reference frequency Plots (RF Plots) were constructed to show the rate for the chosen reference frequency (S or D) as a function of its distance from the best frequency for each tuning curve.

**Table S4.**
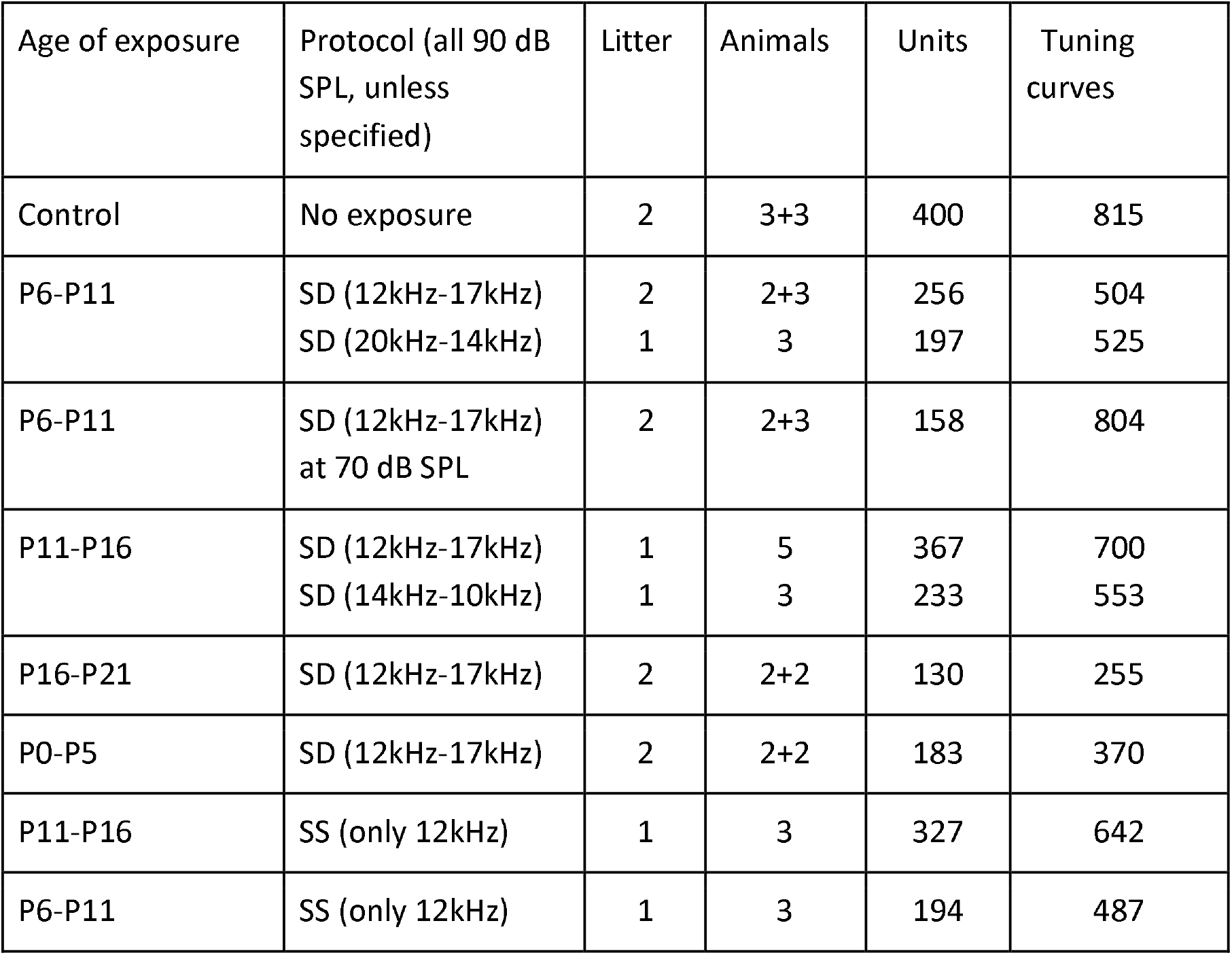

| Age of exposure | Protocol (all 90 dB SPL, unless specified) | Litter | Animals | Units | Tuning curves |
| --- | --- | --- | --- | --- | --- |
| Control | No exposure | 2 | 3+3 | 400 | 815 |
| P6-P11 | SD (12kHz-17kHz) | 2 | 2+3 | 256 | 504 |
|  | SD (20kHz-14kHz) | 1 | 3 | 197 | 525 |
| P6-P11 | SD (12kHz-17kHz) at 70 dB SPL | 2 | 2+3 | 158 | 804 |
| P11-P16 | SD (12kHz-17kHz) | 1 | 5 | 367 | 700 |
|  | SD (14kHz-10kHz) | 1 | 3 | 233 | 553 |
| P16-P21 | SD (12kHz-17kHz) | 2 | 2+2 | 130 | 255 |
| P0-P5 | SD (12kHz-17kHz) | 2 | 2+2 | 183 | 370 |
| P11-P16 | SS (only 12kHz) | 1 | 3 | 327 | 642 |
| P6-P11 | SS (only 12kHz) | 1 | 3 | 194 | 487 |

For example, if the reference frequency (RF) is chosen to be 12kHz (S), and for a given tuning curve with the best frequency at 24kHz, the rate at RF is 12kHz is 10sp/s, then the RF Plot will have a value of 10sp/s 4 bins (1 octave) to the left of centre for that tuning curve. This is done for all the tuning curves, and the population RF curve was Plotted. The grey bar for the control was constructed by repeating the same process 13 times considering each frequency as the reference frequency and finding the maximum value of the rates at each bin, thus signifying the maximum boundary. The bar Plots were constructed by normalizing the RF rates at the best frequency (centre of the RF curve). The normalization was done by dividing it with the mean of all the BF rates except the ones within a distance of a half octave on both the sides of the RF curve.

#### Connectivity between pairs (Fig. 7)

Cross-correlation between pairs of PSTHs (nrep= 20-30 repetitions of SD stimulus and swap, bin size = 5ms) was used to detect connections between simultaneously recorded pairs of neurons (SP-L4 pairs). Bootstrap analysis was done to get variability in the cross-correlograms (MATLAB ‘xcorr’ function, up to lag of 20 ms) by randomly resampling (same number of repeats from the existing 20-30 repeats) with replacement, 100 times. Thus a mean and SD (dark line with errorbar (Fig. 7B) of the correlograms were obtained. To compare with spurious cross-correlations, all repeats from both neurons were mixed to give 2*nrep (number of repetitions) spike trains, and randomly a number of reps of them were assigned to each of the neurons, and from these PSTHs, cross-correlograms were obtained with mean and SD by repeating the process 100 times (gray line with errorbar, Fig. 7B). Non-overlap of errorbars were considered to be significant connections between the pair of neurons. We used 1.65-SD for data in P12 and above groups and 1-SD in before ECO group due to the large difference in firing rates of single units in the two cases.

The cross-correlation at best provides a linear time-shifted dependence between the activities of L4 and SP, although convincing enough for our result, it is unable to capture the changing temporal co-variability between the two signals. To address this issue, we also verified the connectivity proportion results using granger causality, which provides more stringent criteria to find the direction of the information flow (causation). We performed this between the mean PSTH in 5ms binning of the SP and L4 responses.

### Details of Binary Network Model of L4 and SP with TC Inputs

We built a binary neural network consisting of two separate thalamic input units for standard and deviant. The thalamic units project to a sub-plate neuron and further to a cortical neuron. The sub-plate neuron also projects to the cortical neuron. A similar model was used to demonstrate ocular dominance in visual cortex^4,5^. The spike rate of the two thalamic inputs are given by,

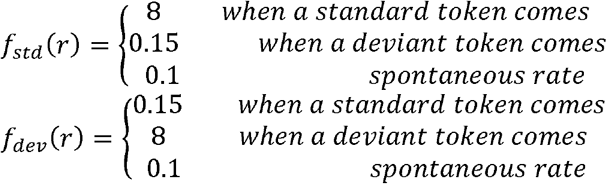

The synaptic weight of the thalamo-subplate projections was kept constant at a value of 0.2. The projections received by the cortical neuron was kept plastic and learning was implemented following the below plasticity rule.

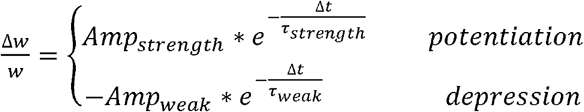

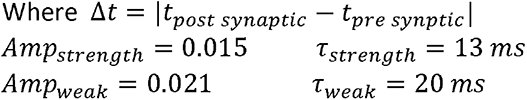

We performed simulations for two regimes, one for the early development phase and one for the later. For the early phase, we kept the synaptic weight of the sub-plate projections at a value of 0.22 and thalamo-cortical projections at a smaller value of 0.02. It was done to mimic the early stages of cortical development where the thalamic projections are unable to drive the cortical activity alone and are aided by the sub-plate for driving the cortical activity ^1,2,6^.

In the later phase, both these weights were kept at an intermediate value of 0.04 and 0.12 respectively. This reflects the intermediate stage of development where the thalamo-cotical projections are strengthening, and the sub-plate-cortical connections are weakening^2,6^.

All the synapses were kept depressing throughout the simulation with the following equations governing their dynamics^7^,

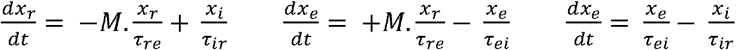

Where M=1 whenever a spike occurs.

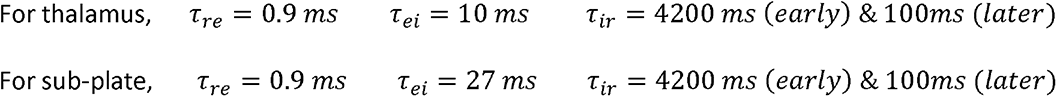

The value of *τ_ir_* was chosen in accordance with our electrophysiology data which shows a decrease in adaptation in later ages.

The sub-plate and the cortical neuron were modelled on the principles of integrate and fire neuron^1^. The membrane potential can be written as follows,

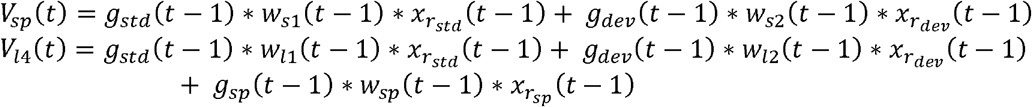

Where g(t) is the conductance and can be written as,

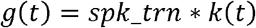

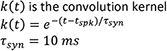

The threshold of Layer IV neuron was kept constant at 0.1 for both the regimes, while that of the subplate neuron was kept at 0.05 for the early regime and 0.15 for the late regime.

The refractory period dynamics after each spike was modelled with a separate exponential given by^1^,

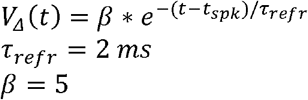

The voltage drops by the above value after every spike. The refractory period lasts for 20ms after the spike. A leak of 10% was also incorporated^1^.

The model gets a drive from the thalamic activity, modelled by uncorrelated Poisson spikes in the two thalamic units. The stimulus train consisted of standard and deviant tokens occurring with a probability of 0.9 and 0.1, respectively in the standard-deviant protocol and consisted of only standard tokens in case of only standard protocol. Each token is 50ms long separated by a 250ms inter-token-interval with a sampling rate of 1000/sec.

### Theoretical Analysis for Optimization based on Mutual Information Maximization and Sparseness

We consider the same stimulus train that we used in the exposure experiments.

We assume the probability of deviant to be x and that of a standard to be 1-x

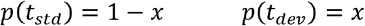

The responses for each of the two tokens are normally distributed

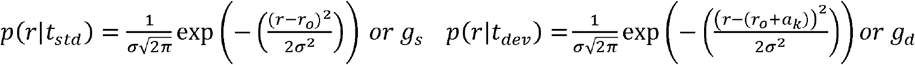

We consider the variance *σ*^2^ and mean response for the standard token *r*_0_ to be constant. The mean response for the deviant token varies with each successive token, and its difference from mean standard response takes values *a_k_* for every kth token. We assume all the tokes to be independent and without any history dependence (no adaptation). Thus,

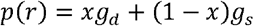

We wish to know the trend of *f*(*k*) with successive tokens that will enhance the discriminability between the standard and the deviant token. To do that, we use mutual information as a measure of discriminability paired with a sparse coding constraint and try to maximize it.

Our constrained information for the k^th^ token is,

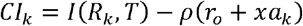

Where *I*(*R_k_, T*) denotes the mutual information between the response at the kth token and the stimulus, *ρ* is the langrage multiplier which is constraining the mean response at the kth token. The lagrangian has been constructed to minimize the mean response at the kth token (sparse representation).

We need to find the value of *p*, which will maximize the mutual information between the entire stimulus train and the total response along with maintaining a sparse representation throughout the stimulus period. We can write the total constrained information as the sum of the constrained information of independent individual tokens,

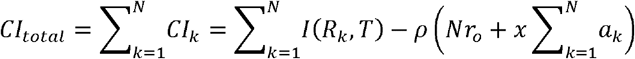

Where N is the total number of tokens.

For any token, *I*(*R_k_, T*) can be written as,

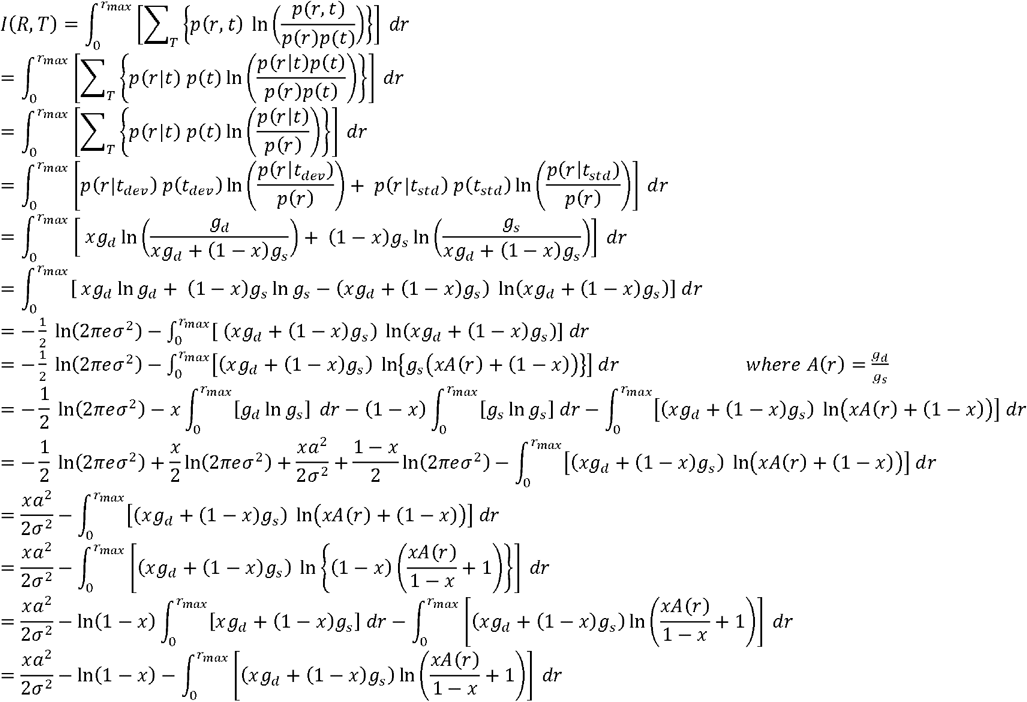

We wish to use the Taylor expansion of ln in the above expression. To do that we need,

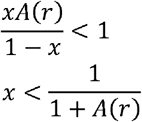

We see that *A*(*r*) is the ratio of two normal distributions and it can be reduced to,

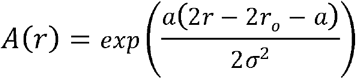

Since *x* is a constant and is only dependent on the stimulus, we set a limit for x that will make this expression valid for all the possible rates,

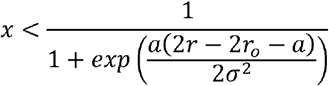

If we assume that on the 1^st^ token (when the discrimination has not yet started), the mean rate for standard and deviant is the same (a=0), we see that,

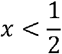

Thus we proceed further with an assumption that the probability of the arrival of the deviant is less than 50% and the results that we discuss later will hold true only on this assumption. We now perform the Taylor series expansion,

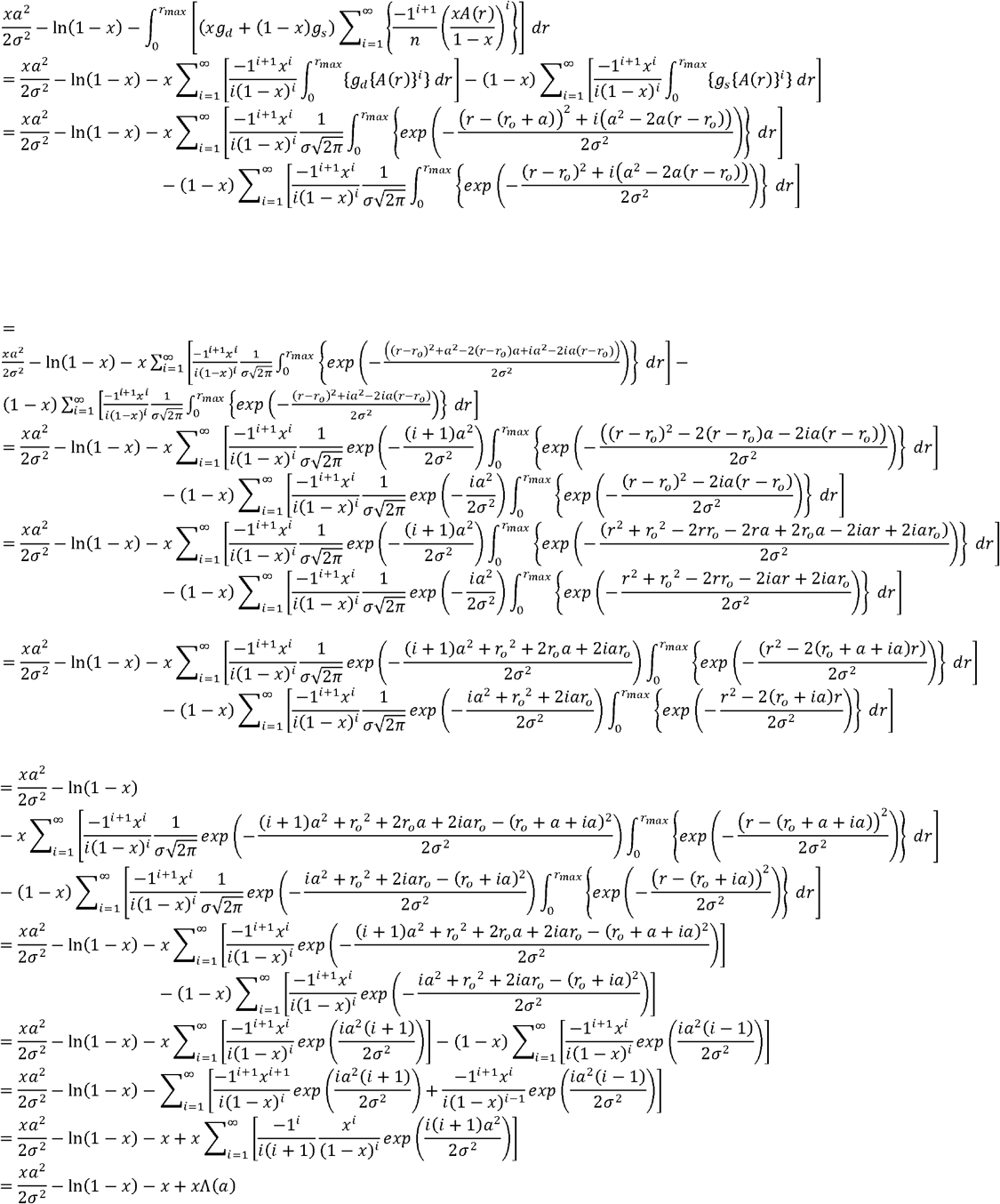

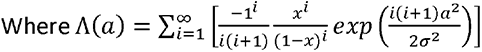

On a careful examination, we see that the above expression will always be positive. We find the change in constrained information between two tokens m&n (m<n)

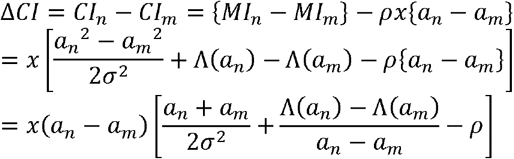

We want the above expression to be greater than or equal to zero. Thus, we get two cases,

Case 1: *a_n_* > *a_m_*

We need,

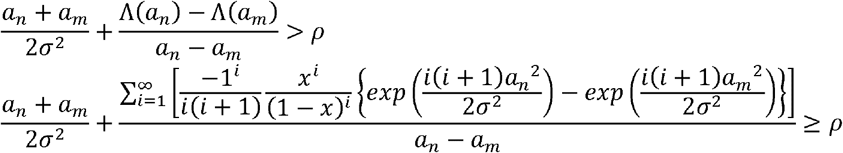

We see that as *a* increases, the 2^nd^ term in the above expression increases with the exponential of the square of the increase in a. The 2^nd^ term overall will be much larger than 1. If we assume a monotonic behavior in *a*, the least value of Λ(*a_n_*) − Λ(*a_m_*) will be between the first two tokens. Thus we can say that,

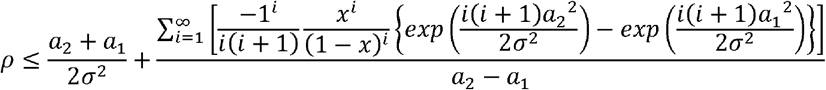

The above expression sets a limit on the value of *ρ* or in other words, the weightage of sparseness which will keep the mutual information increasing.

This case corresponds to the early age results, where the responses for deviant was greater than that of standard (a>0).

Case 2: *a_n_* < *a_m_*

We need,

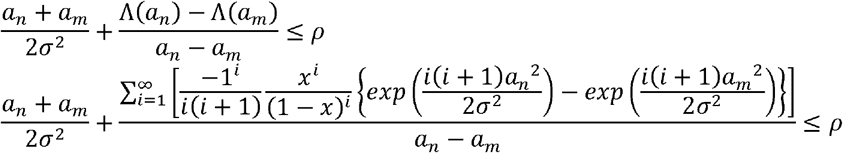

We see that as *a* decreases, the 2^nd^ term in the above expression decreases with the exponential of the square of the increase in a. The 2^nd^ term overall will be much less than −1. If we assume a monotonic behavior in *a*, the largest value of Λ(*a_n_*) − Λ(*a_m_*), will be between the first two tokens. Thus we can say that,

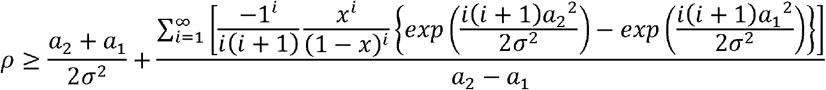

The above expression sets a limit on the value of *ρ* or in other words, the weightage of sparseness which will keep the mutual information increasing.

This case corresponds to the later age results, where the responses for the standard was greater than that of deviant (a<0).

### Supplementary Figures

**Figure S1.**
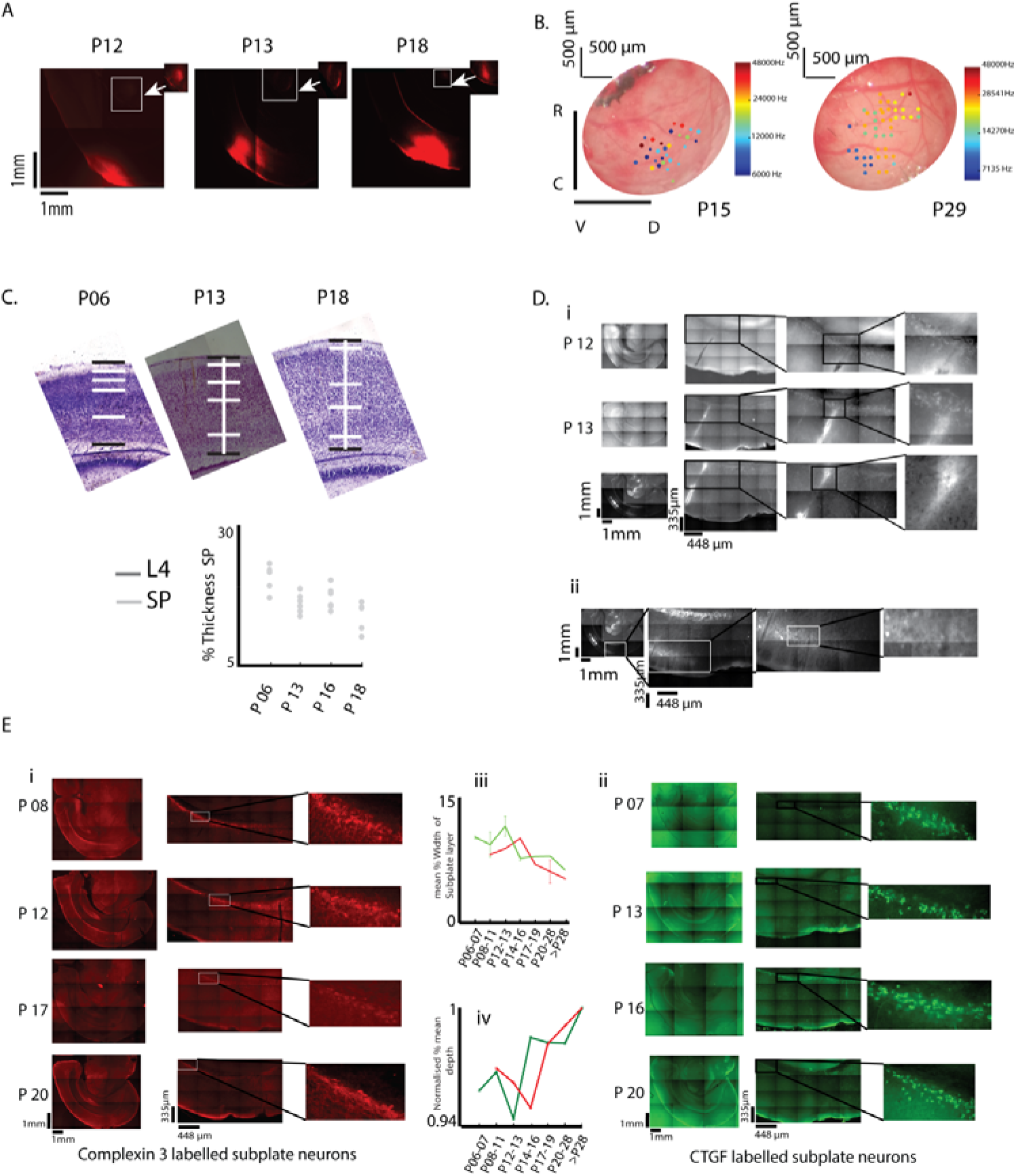
A) Representative confirmation of recording in A1 in lower age groups P12, P13, P18 with DiI injections in the recording site with labelling in MGBv (inset). B) Sample locations of recordings of single units mapped onto the cortical surface showing broad clearer tonotopy in P29 than in P15. C) Thickness of SP as a percentage of cortical thickness over age determined from cell morphology with Nissl stains (Above). D) Slices were cut from brains (fixed) in which single-unit recordings were performed in the ACX (representative examples at 3 ages, P12-P14, Above) and IHC was performed. Electrode tracks show recordings performed in SPN and in case of dual recordings (second electrode type, Fig. 3A) in SCNN1-td-tomato mouse (P14) show electrodes in L4 and SPN simultaneously.

**Figure S2.**
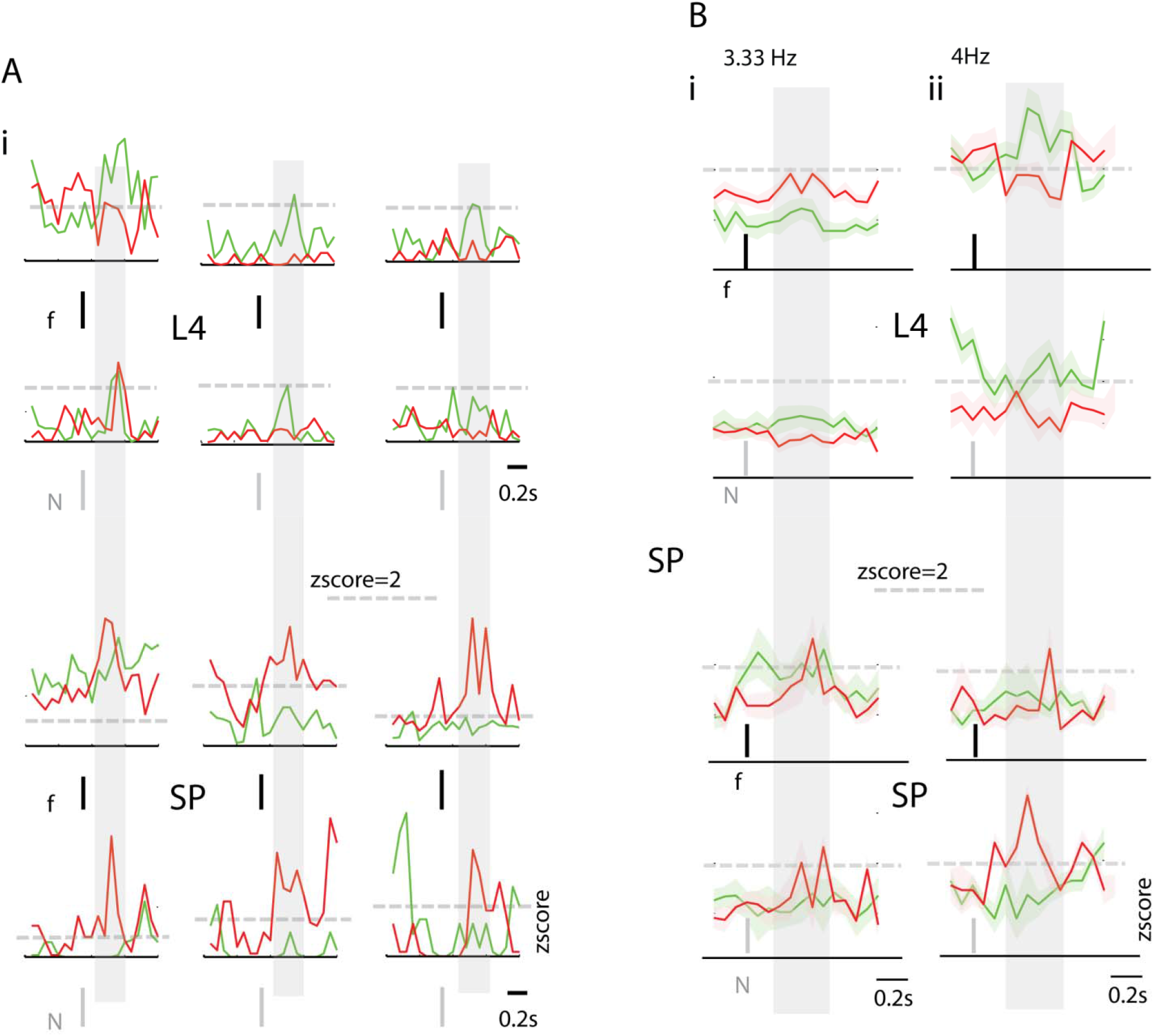
A) i) Example traces of zscores of 3 L4 neurons (upper 2 rows) and 3 SP neurons (lower 2 rows) in the first 3 columns, showing response to tone (f, Standard in Green and Deviant in Red, Above) and noise (N, Standard in Green and Deviant in Red, Below). A light gray patch is the mean rate window (250 ms) across which CSI(F) was computed. B) i) Mean population zscore of all L4 and SP neurons (upper and lower two Plots respectively) in response to tone (f, Standard in Green and Deviant in Red, Above) and noise (Standard in Green and Deviant in Red, below) presented at 3.33 Hz. B) ii) Same as B) i), mean population zscore of L4 and SP neurons in responses to tone and noise, delivered at 4Hz.

**Figure S3.**
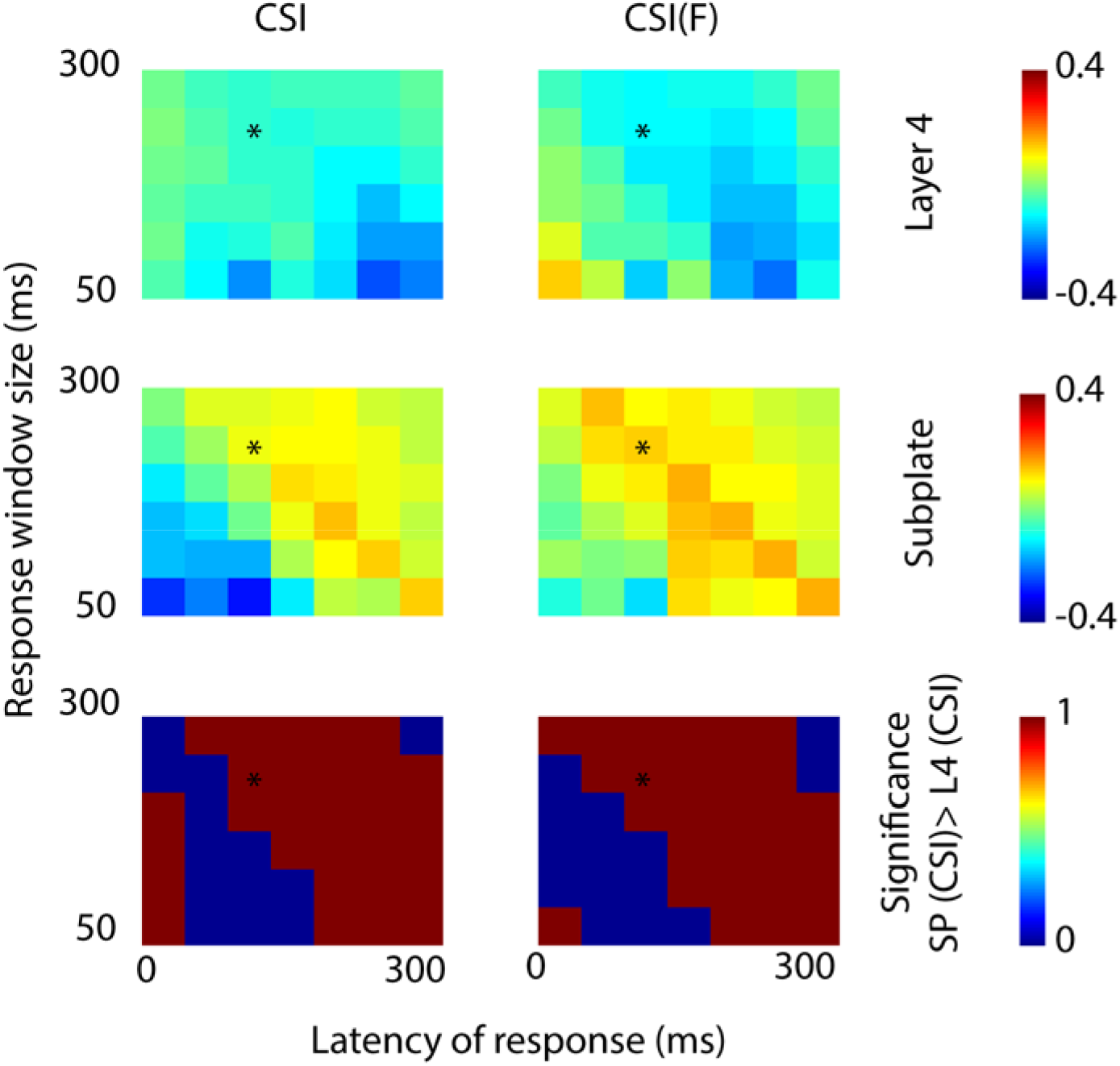
Top and middle row shows the mean CSI values of the population of L4 and SP neurons respectively for different choice of response latencies and response window sizes. Asterisks mark the response latency and response window size combination used in Fig. 4 CSI calculations. The two columns are for the 2 different CSIs used CSI and CSI(F). Bottom row shows which combinations of response attribute have significantly higher CSI in SP than in L4 (brown, blue indicates NS).

**Figure S4.**
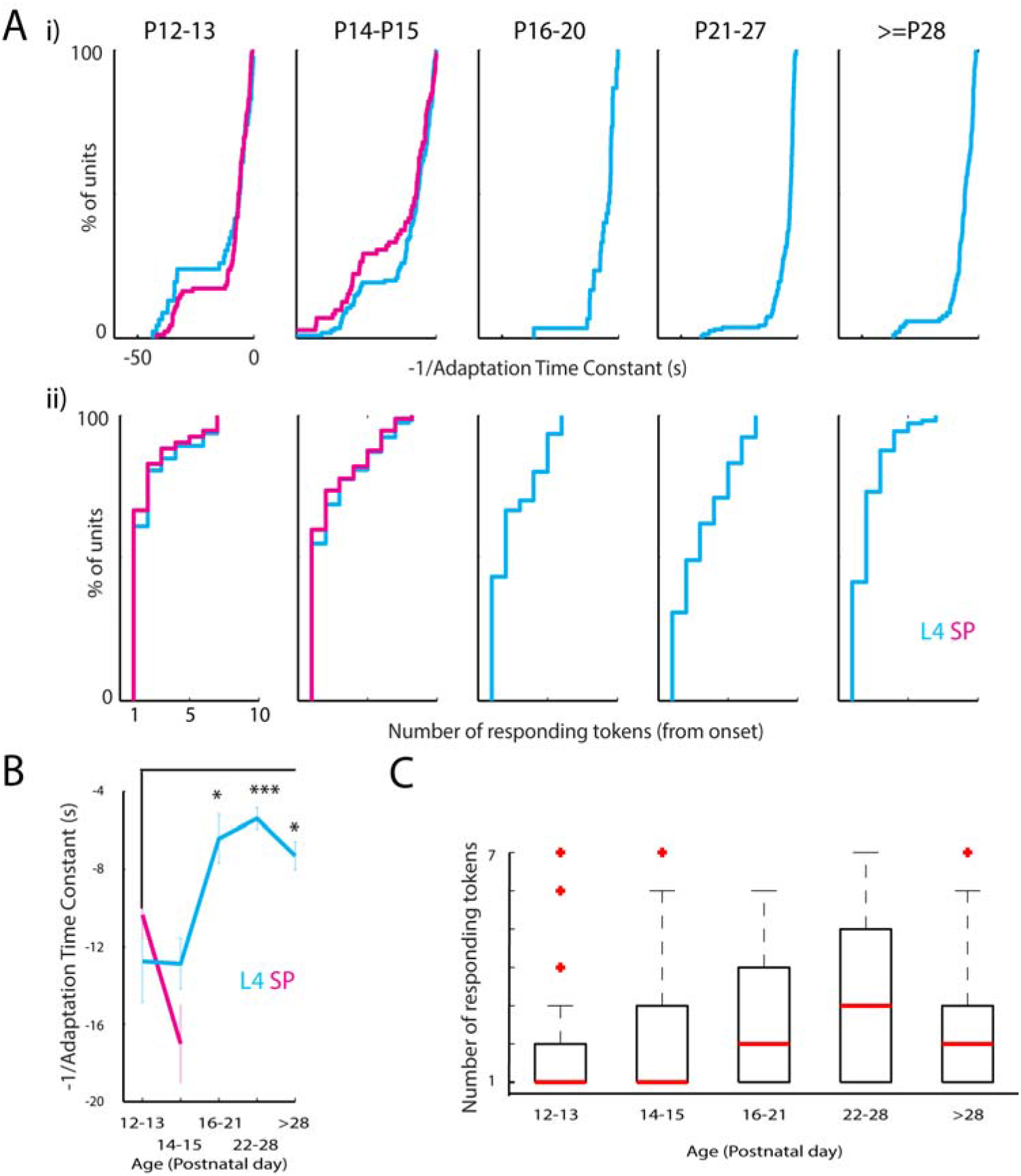
Ai) Population CDFs of adaptation time constants (as −1/time constant) for L4 neurons (Cyan) and SP neurons (Magenta) across different age groups (SP only for P12-15). Adaptation time constants, determined from exponential fits to mean responses to each successive token of Standards, increase with age. Aii) Shows population CDFs (for the same set of neurons and age groups as in the top row) of the number tokens in which there were significant responses in the first set of 7 (or 6) tokens before the deviant for the same population of neurons. B) Mean −1/Adaptation time constants for the data in Ai. Adaptation time constant increases with age (ANOVA, p<0.0001). C) BoxPlots showing range with median, IQR and outliers for data in Aii. Significant responding tokens increases with age (ANOVA, p<0.0001).

**Figure S5.**
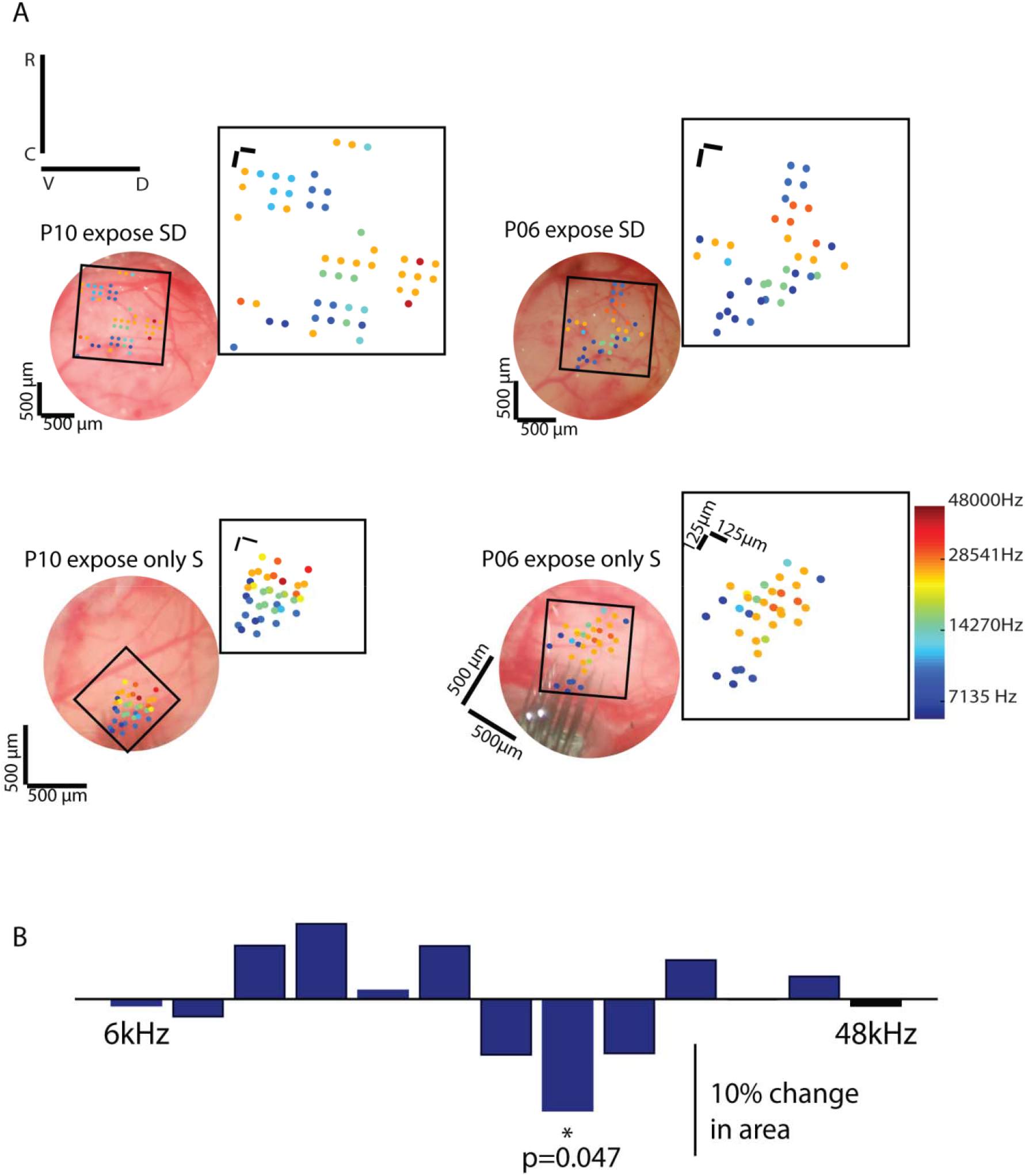
A) Four example images of location (along the cortical surface) of BFs on the cortical surface in the craniotomy of single units recorded in L4 at ages P28 and above (Fig. 6A single-unit recording) identified in color (colorbar, bottom right) for four exposure groups EX2 and EX1 in the top row (SD exposure) and EX5 and EX6 in the bottom row (SS exposure). Broad tonotopy showing A1 and AAF are clear in the images. B) Bar Plot shows the % change in an area of each of the preferred frequencies in A1 and AAF between control (EX0 group of mice) and the only standard P10-P15 group (EX6).

**Figure S6.**
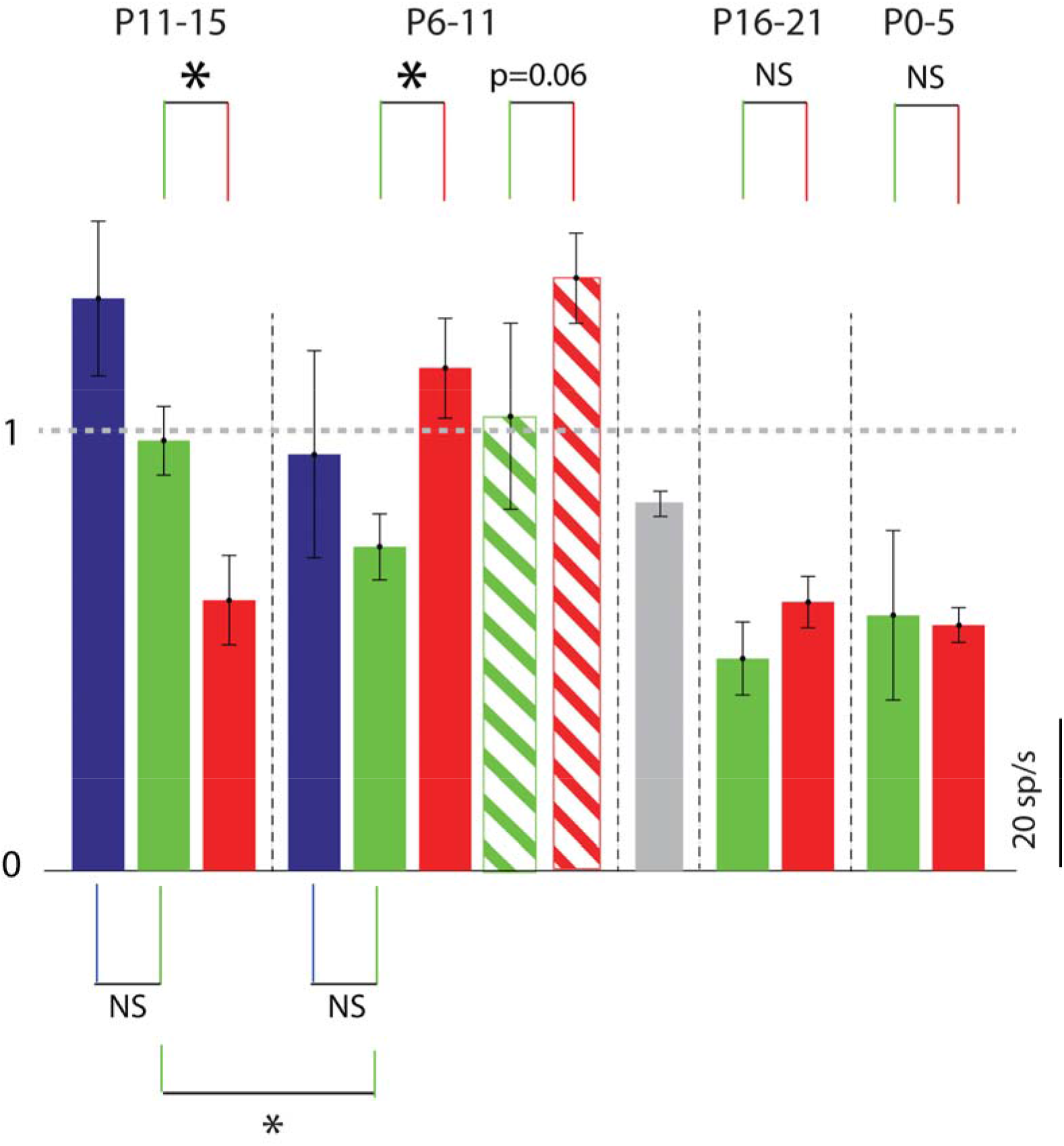
Bar Plot summarizes population mean absolute response to RF for neurons with BF at RF of all 6 EX cases (2 bars each, red and green for SD exposure cases and 1 blue for SS). Mean of the 13 control cases shown in gray bar in the centre. The stripped bar here represents the p6-P10 70 dB SPL exposure protocol (EX5).

### Supplementary Videos

**Supplementary Video 1:** This video demonstrates the widefield imaging-based sound-evoked responses to tone (17kHz, 20dB attenuation, ~ 70 dB SPL) in ACX of an awake Thy1-Gcamp6f mouse at P9. Video acquired at 10 fps.

**Supplementary Video 2:** This video demonstrates the 2-Photon imaging-based sound-evoked responses to tone (24kHz, 0dB attenuation, ~90 dB SPL) in ACX of a Thy1-Gcamp6f mouse at P7. Video acquired at 5 fps.

**Videos are available at https://drive.google.com/drive/folders/1UvD6NXDpHhnKrEB1N2zaoAW4S-gI7jDi**

